# Compositional control of ageing kinetics in TDP-43 condensates

**DOI:** 10.1101/2025.02.21.639421

**Authors:** Nuria H. Espejo, Alejandro Feito, Ignacio Sanchez-Burgos, Adiran Garaizar, Maria M. Conde, Antonio Rey, Alejandro Castro, Rosana Collepardo-Guevara, Andres R. Tejedor, Jorge R. Espinosa

## Abstract

Biomolecular compartments orchestrate the spatiotemporal organisation of cells. The spontaneous assembly of proteins and nucleic acids through liquid-liquid phase separation into biomolecular condensates has been shown to ubiquitously contribute to the functional compartmentalisation of the cytoplasm and nucleoplasm. However, some condensates can undergo an additional phase transition from functional liquid states to pathological solid-like assemblies (i.e., ageing). This liquid-to-solid transition, driven by the accumulation of protein cross-*β*-sheet structures, represents a hallmark of multiple neurodegenerative disorders. In this study, we employ Molecular Dynamics simulations to explore the role of various biomolecules in regulating the ageing kinetics of condensates scaffolded by TDP-43, a key RNA-binding protein linked to amyotrophic lateral sclerosis and frontotemporal dementia. We find that the recruitment of arginine-rich peptides, such as those produced by the C9orf72 gene, accelerates the nucleation of cross-*β*-sheet structures. In contrast, the inclusion of poly-Uridine RNA and the HSP70 chaperone significantly slows the emergence of these structures. Remarkably, we observe a correlation between the compactness of the low-complexity domain of TDP-43— which drives the transition to cross-*β*-sheet structures—and the condensate ageing kinetics as we vary the composition of the condensates. Moreover, we find that near-interfacial regions of TDP-43 condensates exhibit faster *β*-sheet transitions than the bulk core of the condensate. Together, our findings underscore the critical role of client biomolecules in modulating protein conformational ensembles and intermolecular interactions, thereby controlling the propensity of condensates to transition into harmful solid-like states.

## I. INTRODUCTION

Biological function relies on the precise spatiotemporal compartmentalisation of the cell material. While some of the organelles possess membranes which delimit their material from the surrounding medium, such as the cell nucleus^1^, mitochondria^2^ or lysosomes^3^, it has been shown that many other intracellular compartments are membraneless, instead formed via liquid-liquid phase separation (LLPS)^4–9^. LLPS enables the emergence of dynamic compartments—known as biomolecular condensates—which are essential for undertaking numerous cellular functions such as gene expression^10^, cell signaling^6^, or regulation of biochemical reactions^11^, among many others^12,13^. The biomolecular building blocks behind intracellular LLPS are mainly proteins, usually with intrinsically disordered regions (IDRs)^14–17^, and nucleic acids that can establish multiple homotypic and/or heterotypic interactions with other cognate biomolecules, i.e. different IDRs, RNAs, or DNA^18–20^. Whilst initially, the liquid-like behaviour of the molecules within the condensates was thought to be a defining feature of such systems, it has been recently shown that the material properties of biomolecular condensates can encompass low to high viscous fluids, hydrogels, soft-glasses, and even solid-like states^21–23^.

In the last decade, intracellular LLPS has gained significant attention due to its strong implications in physiology and disease^6,24,25^. Dynamic liquid-like condensates can gradually age, even in the absence of changes in their chemical composition and thermodynamic conditions, and gradually transform into kinetically trapped states, displaying a wide range of solid-like material properties^21,26^. Whilst the liquid-like behaviour of condensates underpins their functions during health, kinetically arrested states have been associated with the proliferation of neurodegenerative and age-related disorders^27^. Numerous RNA-binding proteins (RBPs) contain low-complexity domains (LCDs) with high aromatic content^21,28^, which are important contributors to protein multivalency, i.e., the inter-molecular connectivity that enables condensate stability. However, these regions can also lead to disorder-to-order structural transitions, giving rise to inter-protein *β*-sheet structures^29,30^ and amyloid fibrils^27,31^. The fibrillization and ageing of protein condensates with LCDs has been shown to be underpinned by low-complexity aromatic-rich kinked segments (LARKS), which are prone to undergo transitions into cross-*β*-sheets^23,29,30^. Critically, hundreds of RBPs containing LARKS have been identified within the human proteome and are important components of fundamental intracellular structures such as stress granules, ribonucleoprotein granules, P bodies, or germ granules^32–34^. Hence, liquid-like condensates involved in essential cellular functions can also serve as precursors to aberrant solid-like states that no longer provide the necessary environment for intracellular chemical reactions. Relevant examples of protein condensates susceptible to developing ageing are those formed by RBPs such as hnRNPA1^35^, hnRNPA2/B1^6^, Fused in Sarcoma (FUS)^22,36^, or the TAR DNA-Binding protein of 43 kDa (TDP-43)^25^. These different RBPs have been associated with multiple neurodegenerative disorders such as amyotrophic lateral sclerosis (ALS), frontotemporal dementia (FTD), or Alzheimer’s disease, among others^27,31,37–45^.

TDP-43 is a key protein involved in RNA metabolism, transcriptional regulation, and stress responses^24^, which is capable of undergoing phase separation and subsequent condensate solidification^29,46^. Physiologically, TDP-43 is responsible for numerous heterotypic interactions with different biomolecules such as RNAs or DNAs over their interactions with the solvent, and can guide the formation and degradation of membraneless assemblies, as well as contribute to other multiple cellular processes^47^. However, under pathological conditions, TDP-43 is mislocalized in the cytoplasm and sequesters RNA, thus disrupting RNA metabolism^45^. The partly helical region in TDP-43 LCD—which contributes to its propensity to aggregate under pathological conditions^48,49^—harbours almost all of the TARDBP gene mutations which have been identified in different neurodegenerative disorders^30,45^. Accumulation of TDP-43 toxic aggregates impedes normal cellular function contributing to neuronal decline and cell death^42^. Importantly, different biomolecules such as polypeptide repeats^50–53^ or RNAs^54**?**, 55^ can strongly modulate the phase behaviour of TDP-43 condensates. Arginine-rich dipeptide repeat (DPR) proteins, such as poly-GR and poly-PR, are abundant in patients with ALS or FTD and have been revealed to promote phase separation and insolubility of TDP-43 *in-vitro*^50,56–58^ and *in-vivo*^52,59^. In contrast, specific RNA sequences (e.g., poly-Uridine) have been identified to bind to TDP-43, reducing its accumulation in the cytoplasm and potentially stabilizing its liquid-like state preventing harmful aggregation^54,60,61^. Furthermore, chaperones, which often display a high binding affinity to proteins that scaffold biomolecular condensates, promote dynamic regulation of condensate stability, including modulation of their saturation concentration (C_*sat*_), and inhibition of pathological phase transitions^62,63^. For instance, the Heat Shock Protein 70 (HSP70) plays a critical role in disassembling aged stress granules containing misfolded proteins such as SOD1^64^ or TDP-43^65^. Hence, whilst it is known that diverse biomolecules can substantially regulate the phase behaviour of RBPs such as TDP-43, the precise mechanism at the molecular level by which each individual species dictates the material properties, internal organization, and stability of protein condensates remains elusive.

On a macroscopic level, ageing of biomolecular condensates is characterized by the loss of fluidity, condensate reduced fusion, and longer recovery times after photobleaching^12,31,66–69^. Both active and passive microrheology techniques—involving bead-tracking and optical tweezers^70–74^—as well as fluorescence recovery after photobleaching (FRAP) or green fluorescent protein (GFP),^32,70,75^ have demonstrated the continuous drift that initially liquid-like protein^76,77^ (or RNA^78^) condensates experience over time to become kinetically arrested assemblies. However, traditional experimental techniques often struggle to elucidate the molecular intricacies of binding/unbinding events and intermolecular forces that drive condensates out of function^79–81^. Computer simulations, either atomistic^82–87^ or coarse-grained^88–94^, offer a powerful framework to reach submolecular resolution and characterize the interactions and dynamical ensembles of the individual constituent molecules within condensates, as well as the driving forces underlying aberrant aggregation^91,95–100^. Molecular Dynamics simulations of different resolutions have focused on the processes of amyloid formation and cross-*β*-sheet stacking of small peptides and aromatic-rich domains^101–103^. However, less work has been devoted to uncovering the molecular mechanisms of liquid-to-solid transitions in higher-order assemblies such as multicomponent biomolecular condensates as those found in the cell^96,104^. Therefore, there is an urgent need to understand the different governing parameters that control ageing to propose effective strategies to liquefy biomolecular condensates and prevent the proliferation of solid-like assemblies associated with neurodegenerative pathologies.

In this work, we investigate the phase behaviour of multicomponent condensates where TDP-43 acts as the main LLPS scaffold. We examine how the stability, internal architecture and ageing kinetics of TDP-43 condensates are regulated by the inclusion of three types of biomolecules: arginine-rich polypeptides, single-stranded disordered poly-Uridine RNA, and the HSP70 chaperone. We use the sequence-dependent Mpipi-Recharged model^105^, a residue-resolution coarse-grained model that outperforms previous existing force fields for biomolecular phase separation and drastically improves the description of charge effects in condensates^87,105–107^ while still considering implicit solvation for computational efficiency. We benchmark the performance of the model to describe single-molecule properties of TDP-43, as well as higher-order self-assembly of pure TDP-43 condensates and binary mixtures of arginine-rich polypeptides with TDP-43. After validating the model for these systems, we combine equilibrium and non-equilibrium simulations of TDP-43 multicomponent condensates to elucidate the impact of the aforementioned biomolecules in regulating the propensity of TDP-43 to form interprotein *β*-sheet structures. We find that while arginine-rich polypeptides increase the nucleation rate of cross-*β*-sheet nuclei—as experimentally found^52,108^—the recruitment of poly-Uridine RNA and HSP70 substantially decelerate the progressive formation of cross-*β*-sheet clusters. Interestingly, we also detect that inter-protein *β*-sheet transitions occur faster in condensates coexisting with the diluted protein phase, which displays an interface, than in the bulk of the condensate. This behaviour is ascribed to the relative ratio of inter vs. intramolecular contacts that aggregation-prone domains (i.e., LARKS across the LCD in TDP-43) establish depending on the precise condensate biomolecular region and surrounding physicochemical environment. Taken together, our study links the impact of different species on TDP-43 intermolecular contacts to the condensate’s propensity to form cross-*β*-sheet clusters and transition into aberrant solid-like states.

## II. RESULTS

### A. A residue-resolution coarse-grained model for TDP-43 phase-separation

We simulate TDP-43 using the Mpipi-Recharged model^105^, a coarse-grained force field in which each amino acid is represented by a single bead with its own chemical identity. The intrinsically disordered regions of TDP-43 are considered fully flexible polymers, in which subsequent amino acids are connected by harmonic bonds, while the globular domains are treated as rigid bodies,^105^ preserving the structures taken from the corresponding Protein Data Bank (PDB). The different residue-residue interactions in the model consist of a combination of electrostatic and hydrophobic interactions^93,112^ implemented through a Yukawa potential and a Wang-Frenkel potential^113^, respectively (further technical details of the model can be found in the Supplementary Material (SM) Section SI). Importantly, the Yukawa potential permits the parameters to be fine-tuned depending on the specific charged residue pair. Such modification with respect to previous models allows the Mpipi-Recharged to consider that, at a coarse-grained level, the asymmetrical description of attraction and repulsion should compensate for the loss of explicit ions and water, as well as for the aggressive mapping of the many charges that an amino acid carries atomistically, reducing it to just one charge centered on its alpha carbon when coarse-grained.

As an initial benchmark for modelling TDP-43, we compare the single-protein radius of gyration (*R*_*g*_) predicted by the Mpipi-Recharged model against atomistic calculations^25^ and *in vitro* experimental values under diluted conditions^109^. The protein architecture of TDP-43 (Fig. 1A) consists of a partially ordered N-terminal domain (NTD and NLS), followed by two well-folded DNA/RNA-binding domains (RRM1 and RRM2) and a C-terminal domain. The C-terminal domain is composed of a low-complexity domain rich in glycine residues and IDRs, along with a conserved region (CR) that forms an *α*-helical structure (Fig. 1A)^45^. A nuclear localisation signal (NLS)^114^ makes it predominantly located in the nucleus, shuttling between the nucleus and the cytoplasm to perform its functions^25,45^. The determination of *R*_*g*_ (see Section SIV in the SM for further details) provides structural insights into the dynamic conformations that the protein can establish depending on its environment (e.g., solvated in an aqueous solution or in a condensed phase). In Fig. 1B, the *R*_*g*_ probability distribution (P(*R*_*g*_)) of TDP-43 under diluted conditions (dashed curve) predicted by the Mpipi-Recharged model is represented along with the mean value of *R*_*g*_ from all-atom calculations^25^ (pink vertical line; Fig. 1B) and *in vitro* measurements^109^ (purple vertical line; Fig. 1B). Impressively, despite the coarse-grained nature of the Mpipi-Recharged model and the complexity of the multi-domain TDP-43 sequence, the mean value of *R*_*g*_ predicted by our model (*R*_*g*_ = 36.2 ± 1 Å) closely reproduces the experimental value of *R*_*g*_ = 41 Å at physiological conditions (i.e. T=298 K and 150 mM of NaCl) and matches the all-atom value (*R*_*g*_ = 35.6 ± 1 Å) within the uncertainty. Furthermore, we have computed the probability distribution of *R*_*g*_ within the condensate at the same conditions (Fig. 1B; red continuous curve). TDP-43 displays larger *R*_*g*_ values in the condensate compared to the diluted phase due to stabilisation of intermolecular contacts among different protein chains in more extended conformations with respect to the more compact conformations that single proteins exhibit in diluted conditions. This result aligns with prior computational^92,115^ and experimental studies^116^, which reported more extended conformations for various IDPs within condensates compared to dilute conditions. Such conformations arise to maximise the intermolecular connectivity, thereby enhancing the enthalpic gain associated with the formation of a condensed liquid network.

**FIG. 1.**
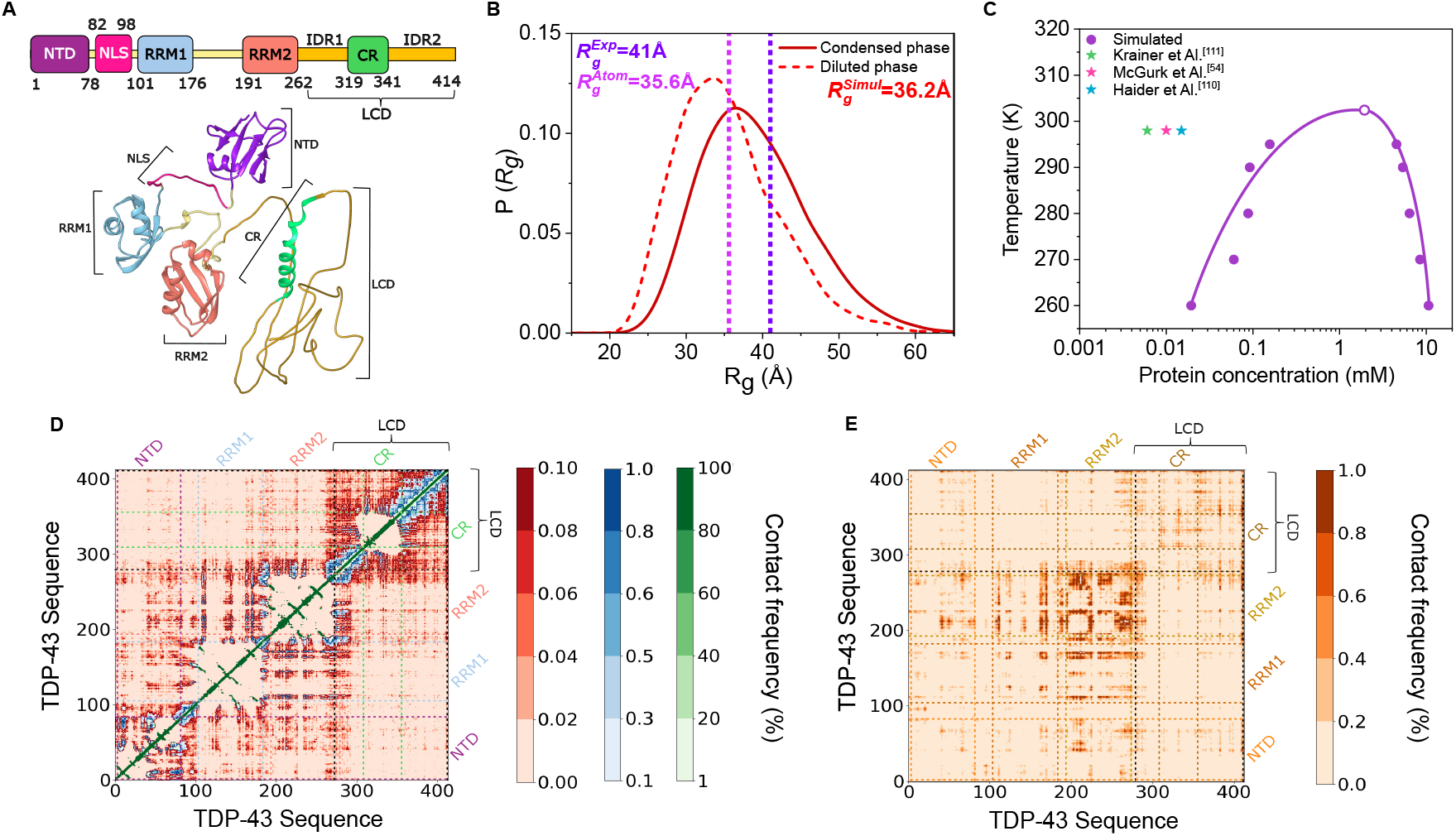
(A) TDP-43 protein sequence broken down in its different domains. It consists of an N-terminal domain (NTD), a nuclear localization signal (NLS), two DNA/RNA binding domains (RRM1 and RRM2) connected by flexible linkers, and a glycine rich C-terminal domain where there is a low-complexity domain (LCD) formed by two intrinsically disordered regions (IDR1 and IDR2) and a conserved region (CR) formed by a *α*-helical structure^25^. The structure of the different globular domains (PDB codes: 5MDI for NTD; 2CQG for RRM1; 1WF0 for RRM2; and 2N2C for CR) across the sequence is shown in the bottom panel (further details on TDP-43 structure are provided in Section SIII of the SM). (B) Probability distribution of the radius of gyration (*R*_*g*_) of TDP-43 within the condensate (continuous line) and in the diluted phase (dashed one) at 300K and physiological salt conditions. The single-molecule mean value of *R*_*g*_ from *in vitro* measurements^109^ and atomistic simulations^25^ at diluted conditions, T=300K and NaCl 150 mM, are depicted by vertical purple and pink dashed lines, respectively. (C) Phase diagram of TDP-43 in the temperature–concentration plane obtained via Direct Coexistence simulations along with the experimental *C*_*sat*_ values calculated experimentally (green, pink and blue stars) from Refs.^54,110,111^. (D) Intramolecular contact frequency map (in %) of TDP-43 at diluted conditions. The different domains are indicated by vertical and horizontal dashed lines. (E) Intermolecular contact frequency map (in %) of TDP-43 within a protein condensate.

To elucidate the precise conditions that enable TDP-43 condensate formation as predicted by the Mpipi-Recharged model, we measure its phase diagram in the temperature–concentration plane (Fig. 1C). The critical solution temperature (*T*_*c*_) for LLPS in TDP-43 is *T*_*c*_ = 302 ± 5 K. Below this temperature, a protein-poor liquid phase coexists with a protein-rich condensed phase. We compare our calculated coexistence line with *in vitro* saturation concentration values (*C*_*sat*_) to undergo LLPS under physiological salt conditions^54,110,111^ (coloured stars in Fig. 1C). The small differences between the reported experimental values can be ascribed to variations in the NaCl concentration (e.g., 50 mM instead of 150 mM in Ref.^111^) and sequence modifications (*C*_*sat*_ in Ref.^110^ refers to TDP-43 LCD, and in Ref.^54^ to a His6-SUMO N-terminally tagged TDP-43-WT). As can be seen, the predicted *C*_*sat*_ overestimates the experimental saturation concentration values by an order of magnitude. Nevertheless, it is important to notice that such discrepancy is acceptable considering that: (1) *C*_*sat*_ can vary as much as few orders of magnitude for protein LLPS depending on the sequence, specific mutations, and the precise thermodynamic conditions^77,117^; and (2) the uncertainty of *C*_*sat*_ evaluated via Direct Coexistence (DC) simulations is relatively high (i.e., up to half-order of magnitude)^106,112^ since the calculation relies on the accurate evaluation of an extremely low protein concentration in the diluted phase that is hardly attainable by the limited time and system size of such simulations. Furthermore, in previous computational studies, we have demonstrated that *C*_*sat*_ cannot be directly compared for certain coarse-grained models (at the same resolution as the Mpipi-Recharged model) with experimental values, since the predicted critical solution temperatures are significantly lower than the melting temperature of pure water^106,107^. Hence, considering all these factors, the difference between the predicted and experimental values (which also involve a certain degree of uncertainty) shown in Fig. 1C seems reasonable.

Once benchmarked the model performance in describing single-molecule properties and higher-order self-assembly of TDP-43 condensates, we elucidate the key molecular contacts that dictate the structural behaviour of TDP-43 at diluted conditions, as well as within phase-separated condensates. To that purpose, we consider a pairwise contact when two residues (either of the same molecule, i.e. intramolecular, or between different proteins, e.g. intermolecular) are found at a distance below 1.2*σ*_*ij*_, being *σ*_*ij*_ the average molecular diameter of the two residues *i* and *j*, and 1.2 a slightly larger distance to the minimum in the interaction potential (∼ 1.12*σ*_*ij*_) between the *i* th and *j* th amino acids (further details on these calculations are provided in Section SVI in the SM). At intramolecular level (Fig. 1D), there are two well-defined regions that exhibit a substantial number of contacts: (i) a large domain corresponding to several globular subregions across the sequence (i.e., NTD, RRM1, and RRM2), and (ii) the most representative segment corresponding to the low-complexity domain (IDR1, CR, and IDR2, please refer to Fig. 1A). The LCD (residues 267-414) is characterized by a repetitive and low-complexity amino acid sequence that favours the formation of intramolecular interactions, and has been suggested to contribute to TDP-43’s propensity to undergo LLPS and subsequent aberrant solidification^30,118–123^. Our simulations show that the LCD tends to self-organize and establish significant interactions within the same TDP-43 molecule (Fig. 1D), largely due to its intrinsically disordered nature and high aromatic composition^115,124^. In a different level, the intermolecular contact map of TDP-43 (Fig. 1E) reveals substantial interaction hotspots (in darker colors) that potentially enable the formation of condensates. The most representative region includes self-interactions of the RRM2 DNA/RNA binding domain of TDP-43 followed by RRM1-RRM2 and LCD-LCD contacts. It has been recently shown that RRM1 partially unfolds in the condensate leading to solvent exposure of cysteine 173 and 175 favouring intermolecular disulfide bond formation, and thereby lowering the minimum concentration of TDP-43 molecules needed to undergo phase-separation^125^. Consistently, in our simulations these two residues display high contact frequencies within the RRM1 region, but even higher with the RRM2 region due to a specific tryptophan in the sequence (Trp-172) which facilitates such strong interconnectivity (please see the protein sequence in Section SIII in the SM). The Trp-172 is one of the TDP-43 exposed tryptophans responsible for the modulation of its toxicity^109,126^. Importantly, although the phase modulation of TDP-43 is primarily supported by RRM1 and RRM2 interactions, the LCD of TDP-43 ranks as the second major domain in terms of establishing the highest intermolecular contact frequency (Fig. 1E). Specifically, there are three tryptophan residues (Trp-334, Trp-385 and Trp-412) involved in condensate stabilization (as reported in Ref.^124^) which form multivalent connections among themselves and with other aromatic and positively charged residues across the sequence (Figs. 1E). Moreover, additional aromatic residues within the LCD, such as Tyr-374, Phe-397, and Phe-401, significantly contribute to LLPS-stabilising interactions through the formation of *π*-*π* contacts. Together, our contact analysis, single-molecule conformational study and phase diagram demonstrate that, despite being a CG model, the Mpipi-Recharged fairly reproduces key experimental features of TDP-43 phase behaviour.

### B. Arginine-rich peptides and near-interfacial condensate regions enhance inter-protein *β*-sheet structural transitions

The propensity of TDP-43 to undergo phase-separation and further *β*-sheet fibril formation has been shown to be strongly regulated by the presence of different biomolecules, including RNA strands of different lengths^61^, or short arginine-rich peptides with varying compositions^50,51,60,127^. To elucidate the key intermolecular forces controlling this behaviour, we first investigate how different peptide di-repeat sequences—GP_25_, GR_25_ and PR_25_—modulate the stability of TDP-43 condensates as a function of their concentration. We define the critical solution temperature for LLPS (*T*_*c*_) as our metric of biomolecular condensate stability^107^ when comparing between different protein/peptide systems. Since imposing a concentration of peptides within a protein condensate is hardly attainable through DC simulations^128^, we perform NpT (constant number of particles (*N*), pressure (*p*) and temperature (*T*)) simulations for protein/peptide mixtures, where *T*_*c*_ is estimated to be between the highest temperature at which the condensate is stable at *p*=0 and the lowest one at which the condensed phase evolves into the diluted phase (further details on this method are provided in Section SII in the SM). This approach—validated in Ref.^128^—additionally allows to evaluate the density of the condensate as a function of temperature for different peptide sequences and/or protein/peptide stoichiometries.

We evaluate the stability of TPD-43 condensates as a function of the GP_25_, GR_25_, and PR_25_ concentration in Fig. 2A. While both GR_25_ and PR_25_ enhance the stability of the condensates up to molar ratios of ∼ 2, the inclusion of GP_25_ consistently hinders condensate formation even at relatively low concentrations (below a molar ratio of 0.25). This is attributed to the strong interactions between the positively-charged arginine-rich peptides (GR_25_ and PR_25_) with the negatively charged and aromatic residues of the TDP-43 sequence. Nevertheless, re-entrant phase behaviour with a maximum in condensate stability can be observed at peptide/protein molar ratios approaching 1:1, driven by peptide-peptide electrostatic self-repulsion occurring at high concentration. This behaviour is consistent with previous experimental findings for complex coacervates of RNA-binding proteins in the presence of charged polypeptides^50,52,108^. Similar non-monotonic changes in the stability of multicomponent condensates have also been reported for mixtures of FUS-LCD and hnRNPA1-LCD^129^, and mixtures of PRC1 and RING1B^130^.

**FIG. 2.**
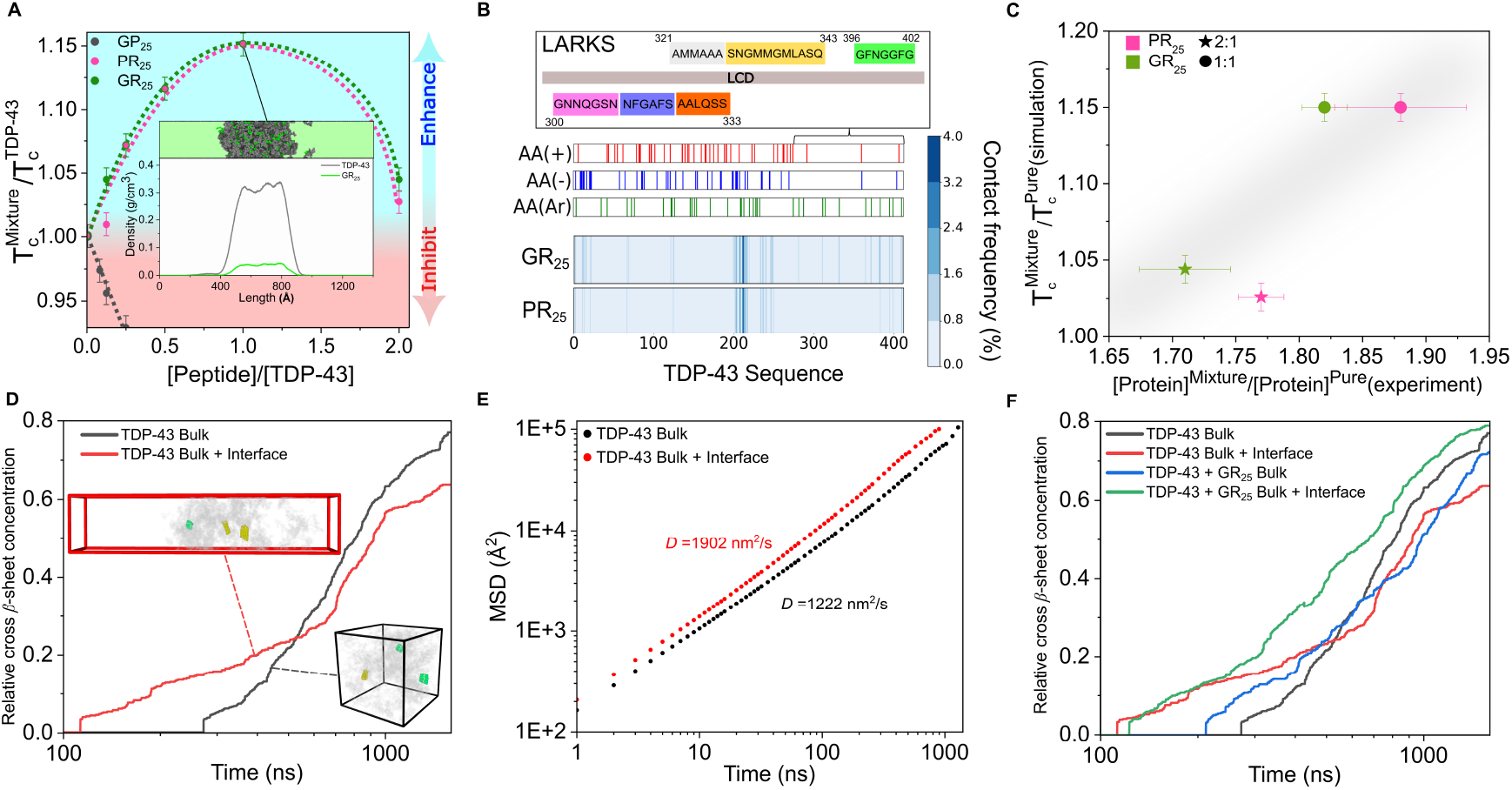
(A) Relative variation in the critical solution temperature as a function of the peptide/TDP-43 molar ratio for different inserted peptides as indicated in the legend. The blue shaded area indicates LLPS enhancement while the red one phase-separation hindrance. Inset: Density profile of a Direct Coexistence simulation of GR_25_ and TDP-43 at 1:1 molar ratio depicting the relative density of each specie across the condensate. (B) Top: Low-complexity aromatic-rich kinked segments (LARKS) located at the TDP-43 LCD. Bottom: Intermolecular contact frequency map (in %) of GR_25_ and PR_25_ with the TDP-43 sequence (414 residues) in each binary mixture at 1:1 molar ratio and 300K. The identity of charged and aromatic residues across the TDP-43 sequence are indicated above. (C) Ratio of the critical solution temperature of the binary mixture over that of pure TDP-43 condensates from Mpipi-Recharged simulations (y-axis) vs. *in vitro*^50^ TDP-43 relative concentration within the condensates in peptide-protein mixtures over pure TDP-43 condensates (x-axis). Each color corresponds to the type of peptide used and the symbol shape corresponds to peptide:protein molar ratio. (D) Time-evolution of the cross-*β*-sheet relative concentration in TDP-43 condensates under bulk conditions (black curve) and within condensates in coexistence with the protein-poor diluted phase (i.e., Direct Coexistence simulation; red curve). Snapshots of both simulation setups are displayed: protein residues forming cross-*β*-sheet clusters are highlighted while the rest of the protein residues within the condensate are faded in grey. (E) Mean square displacement (MSD) of TDP-43 proteins in the systems shown in panel D computed before the emergence of cross-*β*-sheet transitions. The diffusion coefficient (*D*) extracted from this analysis is also included. (F) Time-evolution of the relative cross-*β*-sheet concentration of pure TDP-43 condensates under bulk conditions (black curve), within condensates in coexistence with the protein diluted phase (red curve), and for (1:1) TDP-43:GR_25_ mixtures in bulk conditions (blue) and in coexistence to the diluted phase (green).

Furthermore, complex coacervates consisting of short positively-charged synthetic peptides (SR8 and RP3) and single-stranded RNA poly-Uridine (polyU)^131^, as well as RNA-binding proteins (such as EWSR1, TAF15, hnRNPA1 and FUS) in presence of polyU exhibit an RNA-concentration dependent re-entrant phase transition controlled by the balance between heterotypic associative electrostatic interactions and self-repulsive homotypic electrostatic interactions. In contrast to such reentrant phase behaviour, GP_25_ inhibits the formation of condensates even at low concentrations since glycine and proline, which are both uncharged non-polar residues, can only act as ‘spacers’^117^, and poorly contribute to forming LLPS-stabilising contacts. To further characterise the mechanism by which GR_25_ and PR_25_ modulate TDP-43 phase behaviour, we carry out DC simulations to determine the internal organisation of the different components within the droplets. Strikingly, we find that at the concentration that maximises condensate stability (i.e., molar ratio of 1), arginine-rich peptides remain homogeneously distributed through the condensate core with a marginally low concentration at the interface (Fig. 2A inset). This condensate architecture differs from that previously observed in FUS/polyU mixtures where short molecules (e.g., RNA strands of 50 nucleotides) are predominantly localised at the condensate interface^128^. In this regard, the strength of the electrostatic and cation-*π* interactions between TDP-43 and the arginine-rich peptides underpins their retention in the condensate bulk—to maximise the enthalpic gain and the liquid network intermolecular connectivity—rather than their co-localisation at the interface.

To further understand which specific intermolecular interactions promote the assembly of the condensate upon the recruitment of arginine-rich peptides, we calculate the contact frequency of GR_25_ and PR_25_ peptides with the amino acid sequence of TDP-43 (Fig. 2B). Interestingly, most heterotypic contacts between these peptides and TDP-43 occur through the RRM2 domain due to its strong abundance of negatively charged and aromatic residues. Moreover, the negatively charged residues contained in the NTD and the aromatic residues within the LCD—identified as the key region driving the pathological aggregation of TDP-43^132–137^—contribute to the condensate intermolecular connectivity in presence of arginine-rich peptides. In that respect, recent experimental evidence has shown that arginine-rich peptides bind to regions adjacent to the LCD (e.g. the RGG domains in the FUS protein), enhancing protein aggregation by increasing LCD local concentration and its subsequent inter-protein stacking^138^. A qualitative comparison between our computational predictions for the regulation of TPD-43 LLPS in presence of GR_25_ and PR_25_ peptides against *in vitro* results^50^ is shown in Fig. 2C. Our model correctly predicts the experimental observation that the 2:1 peptide/TDP-43 molar ratio translates into lower phase separation capacity (measured through the relative ratio of *T*_*c,mixture*_*/T*_*c,T DP* −43_ in our simulations and experimentally through the relative protein TDP-43 concentration within the condensates in absence vs. presence of peptides) than the 1:1 stoichiometry. Moreover, the Mpipi-Recharged correctly predicts the similar impact that both GR_25_ and PR_25_ induce on TDP-43 condensate stability (Figs. 2A and 2C). At a 1:1 stoichiometry, both peptides increase protein density within the condensates in both simulations and experiments. This increase is more pronounced compared to stoichiometries in which the peptide concentration is doubled (2:1) or compared to pure TDP-43 condensates. These results reassure the realism of our model in describing the impact of short peptides recruitment with different composition on the phase behaviour of TDP-43 condensates.

To elucidate the impact of arginine-rich peptides in TDP-43 ageing kinetics, we shall now perform non-equilibrium MD simulations incorporating our ageing algorithm, which accounts for the description of structural and energetic transformation of disordered LARKS into cross-*β*-sheet assemblies^23,139^. As input for the free-energy binding difference between the structured vs. disordered LARKS, we use atomistic potential-of-mean-force (PMF) calculations from Refs.^23,82,104,139^. In these studies, the unbinding of cross-*β*-sheet clusters is evaluated based on the structures of Refs.^29,30^, resolved via Cryo-Electron Microscopy (Cryo-EM). PMF calculations are performed using an atom-level force field^140^ which considers both structured conformations from Cryo-EM^29,30^ and intrinsically disordered ensembles from all-atom simulations^140^ to compute the binding free energy difference of the structured vs. disordered state of a cluster composed of 4 to 6 LARKS. Our ageing algorithm evaluates LARKS high-density fluctuations throughout the simulation using a distance criterion based on a local order parameter developed by us^82^. When at least five LARKS from different protein replicas meet specific proximity conditions (i.e., within a cut-off distance consistent with the free energy minima of the PMF calculations), the interactions between these segments are approximated according to the PMF structured binding free energy of the previously studies sequences^23,82,104,139^. Structured cross-*β*-sheet interactions are substantially stronger than LARKS disordered-like interactions (i.e., 3-6 k_*B*_T per LARKS when disordered and 30-50 k_*B*_T upon the structural transition^23^). Furthermore, in addition to the energetic implications of the structural transition, our algorithm also imposes an angular potential between consecutive LARKS residues to account for the increased rigidity of the newly formed structures, thus mimicking the conformational change across the involved segment. The TDP-43 LARKS considered^30^ in our coarse-grained simulations are displayed in Fig. 2B (Top panel). For technical details on the implementation of the ageing algorithm, please see Section SVII of SM.

We measure the concentration of cross-*β*-sheets (normalised by the maximum number of cross-*β*-sheets which can be formed) in pure TDP-43 condensates as a function of time (Fig. 2D). We average 5 different independent trajectories (i.e., with a different initial velocity distribution per trajectory) using two different setups. First, we employ NVT simulations at the equilibrium condensate density at 285K (black curve in Fig. 2D) to emulate bulk condensate conditions below the critical solution temperature. Then, at the same temperature, we perform DC simulations (5 independent trajectories) to represent a condensate that coexists with the protein diluted phase (red curve in Fig. 2D), and thus they account for the presence of an interface in the condensate. Strikingly, we find that the presence of condensate interfaces reduces the nucleation time—i.e., defined as the initial lag time for the formation of the first *β*-sheet nuclei—compared to condensate bulk conditions (Fig. 2D). Whilst we do not observe a substantially higher probability of cross-*β*-sheet formation at the condensate interfacial regions in our DC simulations (when tracking the position of the formed *β*-sheet nuclei), we consistently find that the nucleation rate is 2 times faster in condensates in coexistence with the dilute phase than in those under bulk conditions. Hence, our simulations suggest that interfacial (or near-interfacial) regions in TDP-43 condensates may facilitate high-density local-order fluctuations driving LARKS *β*-sheet structural transitions. Interestingly, these observations for TDP-43 condensates are consistent with previous experimental and computational findings for other RNA-binding proteins such as FUS^76,82^, *α*-synuclein^141^ or hnRNPA1^142,143^, which have been observed to undergo enhanced *β*-sheet fibrillization at the condensate boundaries.

To further explore the origin of such behaviour, we also evaluate the mean-square displacement (MSD) (see Section SVIII in the SM for further details) of the TDP-43 proteins in condensates at bulk conditions and within DC simulations (excluding the marginal number of proteins that for a period of time transitioned from the condensate to the diluted phase; Fig. 2E). We observe a moderately larger diffusion coefficient (*D*) for TDP-43 in condensates with interfacial regions compared to bulk systems, despite similar (though slightly slower at the interface) protein densities. This faster dynamics of TDP-43 in condensates coexisting with the diluted phase may contribute to their faster ageing kinetics. Moreover, in addition to faster protein diffusion, the high-density local fluctuations of the protein LCDs (as recently reported by us for FUS condensates^76,82^) and the more extended LCD conformations—displaying a higher ratio of inter-vs. intramolecular contacts (as discussed in Section II E)—found in condensates coexisting with the diluted phase are likely key contributors to facilitating inter-protein structural transitions that form cross-*β*-sheet clusters.

Next, we analyse the impact of arginine-rich peptide inclusion in TDP-43 ageing kinetics. For that purpose, we measure the time-evolution of cross-*β*-sheet clusters in TDP-43/GR_25_ condensates at 1:1 stoichiometry (Fig. 2F). We choose this concentration since, as shown in Fig. 2A, it maximises the stability of the condensates. Moreover, given that both GR_25_ and PR_25_ establish a similar pattern of intermolecular contacts with TDP-43 (Fig. 2B), and affect similarly its phase behaviour as a function of their relative concentration (Fig. 2A), we only perform simulations for the GR_25_ peptide sequence. We consistently find (for the two sets of 5 different independent trajectories using both bulk NVT condensate simulations and DC simulations) that condensates with interfaces (i.e., DC setup; green curve in Fig. 2F) exhibit faster cross-*β*-sheet nucleation than condensates in bulk conditions (blue curve in Fig. 2F). Furthermore, if we compare pure TDP-43 condensates vs. TDP-43/GR_25_ mixtures under bulk conditions, we observe that the presence of GR_25_ not only promotes phase-separation, but also accelerates cross-*β*-sheet formation. Nevertheless, such effect is negligible in the presence of interfaces where both pure TDP-43 and TDP-43/GR_25_ exhibit remarkably similar nucleation times (red and green curves, respectively). We hypothesise that such different behaviour in bulk conditions can be driven by GR_25_ predominantly interacting with the RRM2 domain of TDP-43, therefore facilitating TDP-43 LCD-LCD intermolecular contacts which in the absence of GR_25_ were compromised in RRM2-LCD interactions (Fig. 1E). However, if LCD-LCD high-density fluctuations occur in near-interfacial regions, the impact of GR_25_ addition in TDP-43 ageing kinetics becomes negligible (as shown in Fig. 2F) as GR_25_ primarily co-localises at the condensate core (Fig. 2A). In agreement with our simulations, arginine-rich peptides have recently been shown to promote neuronal ageing in mice, inhibit ribosome biogenesis, a critical process for protein synthesis, and lead to cellular dysfunction and premature ageing^144^. Hence, arginine-rich peptides not only enhance condensate stability but also increase the ageing rate of protein condensates into kinetically trapped assemblies. Nevertheless, the growth of cross-*β*-sheet structures upon nucleation is roughly similar for all the systems studied in agreement with previous computational^127^ and experimental work^145^.

### C. Poly-Uridine RNA reduces the stability of TDP-43/arginine-rich peptide condensates and their propensity to develop *β*-sheet fibrils

RNA critically regulates the propensity of RNA-binding proteins, such as TDP-43, to undergo phase separation and further maturation^50,128,146–150^. Multiple *in vitro* studies have demonstrated different pathways by which RNAs are capable of modulating protein phase behaviour^61,75,131,151–154^. A particularly relevant example is the RNA-driven re-entrant phase behaviour of numerous protein condensates in which the length, concentration, and sequence of RNA strands tightly regulate the phase diagram and viscoelastic properties of the condensates^61,115,128,131,155^. In this Section, we investigate how polyU single-stranded RNA dictates the stability and ageing kinetics of TDP-43/arginine-rich peptide mixtures. We measure the critical solution temperature (through NpT simulations) of TDP-43 condensates—composed of GR_25_ or PR_25_ and TDP-43 at a 1:1 molar ratio—as a function of the polyU concentration using strands of 250 nucleotides (Fig. 3A). Interestingly, our simulations show that the critical temperature monotonically decreases upon polyU addition (Fig. 3A). To better understand this behaviour, we also perform DC simulations to unveil the architecture of the condensates (Fig. 3B; Top panel). We find that RNA predominantly co-localises at the condensate’s core, strongly interacting with GR_25_ (or PR_25_; Fig. S1), driven by attractive electrostatic interactions between R and U. These results are consistent with previous *in silico* work where RNAs of similar length (i.e., from 200 to 400 nucleotides) were found deep within the condensate’s core of FUS droplets contributing to strengthening the overall connectivity of the liquid network of the system^128^. We also compute the contact frequency distribution of polyU strands with TDP-43 (Fig. 3B: Bottom panel). While both GR_25_ and PR_25_ predominantly interact with the RRM2 domain—rich in aromatic and negatively charged residues (Fig. 2B)— polyU broadly interacts with different segments across the TDP-43 sequence rich in positively charged residues such as arginine and lysine (e.g., scattered from the 45th to the 295th residue; Fig. 3B). However, we note that polyU–TDP-43 contacts are notably weaker than those established by GR_25_ (or PR_25_) with both TDP-43 and polyU (Fig. S1). This suggests that, while RNA interacts with TDP-43, the interaction is much weaker compared to its interaction with arginine-rich peptides, which is primarily driven by electrostatic non-specific interactions. Supporting our simulation findings, recent *in vitro* experiments have also shown how arginine-rich peptides (i.e., GR_20_) establish strong electrostatic interactions between their positively charged side chains and the negatively charged RNA backbone driving condensate formation via complex coacervation^156^.

**FIG. 3.**
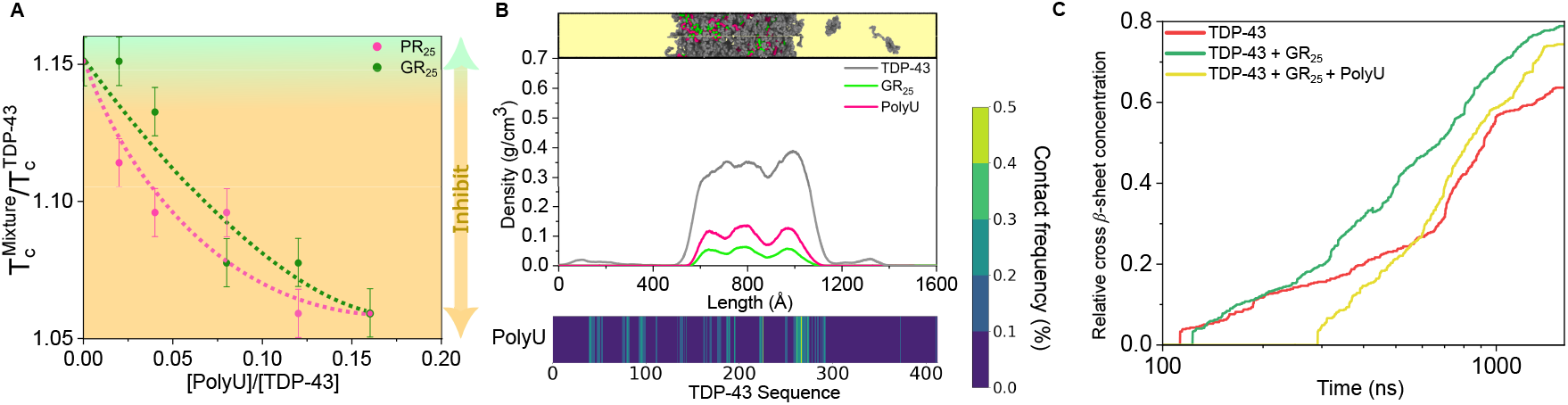
(A) Variation of the critical solution temperature 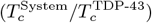 as a function of the polyU/TDP-43 molar ratio in TDP-43/GR_25_ and TDP-43/PR_25_ (1:1 molar ratio) mixtures. (B) Top panel: Simulation snapshot and density profile of a TDP-43/GR_25_/polyU condensate formed by 100, 100, and 16 molecules of each component respectively (i.e., corresponding to a 0.16 polyU/TDP-43 molar ratio) at 285K and physiological salt concentration. Density profiles of each species are shown as indicated in the legend. The *x* axis represents the length of the DC simulation box. Bottom panel: Intermolecular contact frequency of polyU with TDP-43 residues in the condensate shown in Panel B (Top). (C) Time-evolution (averaged over 5 independent trajectories) of the cross-*β*-sheet concentration (normalized by the maximum number of *β*-sheets which can be formed in the system) for TDP-43 (red), TDP-43/GR_25_ (green) and TDP-43/GR_25_/polyU (yellow) condensates in coexistence with the diluted phase.

We further perform non-equilibrium DC simulations (e.g. condensates in coexistence with a diluted phase) to characterise the formation of inter-protein *β*-sheet nuclei in TDP-43/GR_25_/polyU condensates. In Fig. 3C we show the time-evolution of cross-*β*-sheet concentration averaged over 5 independent trajectories with different initial velocities in TDP-43/GR_25_/polyU condensates (yellow curve in Fig. 3C). Inclusion of polyU 250-nucleotide strands (at 0.16 polyU/TDP-43 molar ratio) significantly decelerates (i.e., by a factor of ∼3) the nucleation time of inter-protein *β*-sheet assemblies compared to pure TDP-43 (red curve) and TDP-43/GR_25_ (green curve) condensates. Interestingly, RNA slows down TDP-43-LCD structural transitions without directly establishing frequent polyU–LARKS interactions as shown in Fig. 3B (bottom panel). PolyU outcompetes with TDP-43 to interact with GR_25_, releasing contacts between GR_25_ and the RRM2 domain of TDP-43 (which contribute to enhancing LCD-LCD interactions driving faster cross-*β*-sheet formation). Moreover, direct interactions between polyU and the IDR1 in TDP-43, which is adjacent to several rich-aromatic regions prone to undergo *β*-sheet transitions, are likely to aid in decelerating ageing kinetics. Additionally, RNA-RNA electrostatic self-repulsion at moderately high RNA concentration partially reduces the condensate density decreasing the probability of local stacking of TDP-43-LCD driving *β*-sheet transitions. Overall, the polyU propensity to interact with arginine-rich peptides (Fig. S1) and TDP-43 IDR1 (Fig. 3B) reduces TDP-43-LCD intermolecular contacts (as it will be further discussed in Section II E) and the system kinetics of developing interprotein structural transitions. The addition of single-stranded polyU at high concentrations (i.e., before RNA-driven LLPS inhibition) has also been shown to delay the emergence of cross-*β*-sheet assemblies in other RNA-binding proteins involved in stress granule formation, such as hnRNPA1^139^ and FUS^23^. Moreover, the importance of heterotypic interactions between proteins and nucleic acids and how these interactions influence the mechanistic pathway and kinetics of amyloid formation have been recently highlighted in Ref.^157^. It has been reported how the presence of RNA influences the transition of hnRNPA1A from a soluble state to amyloid, with the pathways being conducted by the RNA/protein ratio: at lower RNA concentration RNA acts as a facilitator for amyloid formation of hnRNPA1A while at higher RNA concentration condensation is inhibited due to reentrant phase behaviour^61,157^.

### D. HSP70 chaperone and RNA decelerate inter-protein *β*-sheet formation in TDP-43 condensates through alternative yet synergistic mechanisms

The heat-shock protein of 70 kiloDaltons (HSP70) was first identified as part of a protective mechanism triggered by heat stress^159^, laying the foundation for understanding its vital role in maintaining protein homeostasis, regulating stress responses, and promoting resilience under adverse conditions^160^. It contains two functional domains: a nucleotide binding domain (NBD) and a substrate binding domain (SBD), which are followed by a C-terminal domain (CTD), as sketched in Fig. 4A (please see the protein sequence in Section SIII in the SM). The two functional domains are connected by a highly conserved inter-domain linker with relevant hydrophobic residues for chaperone function^161^. HSP70 chaperone is one of the most conserved protein sequences throughout different species known in biology^162^, and it acts as an essential regulator of protein folding and cellular stress responses. Furthermore, it has been reported to undergo LLPS *in vitro* and co-phase-separate with different RBPs (such as FUS) preventing them from transitioning abnormally from a liquid state to a solid phase as well as from developing fibrils over time^163,164^. Given its central role in regulating protein phase transitions, this section focuses on exploring its impact on modulating TDP-43/GR_25_ condensates internal organization and ageing kinetics. To that purpose, we first perform DC simulations of a TDP-43/GR_25_/HSP70 condensate at a molar ratio of 1:1:0.16 (Fig. 4B). We note that the structure of HSP70 (shown in Fig. 4A) is maintained across the simulations with the exception of the identified IDRs along the sequence. By performing a density profile of the different species across the condensates, we discover that HSP70 preferentially localises at the condensate’s interface, in contrast to TDP-43 and GR_25_ which are homogeneously distributed throughout the condensate. We note that this is an equilibrium configuration of the condensate since all the species were initially randomly distributed across the simulation box and have since reached equilibrium. To elucidate the key intermolecular interactions triggering such condensate architecture, we compute the contact frequency map of heterotypic interactions between HSP70 and TDP-43 (Fig. 4C). Among the regions of TDP-43 that interact with HSP70, its LCD is one of the most prominent domains in establishing heterotypic contacts, consistent with previous studies^165^ reporting that HSP70 binds to unfolded and predominantly hydrophobic segments of substrate proteins. Additionally, there are two regions of HSP70, within the 250–300 amino acid region—corresponding to a segment across its NBD region—which exhibit high contact frequencies with a significant part of TDP-43 sequence ranging from the 3rd to the 90th residue, and from the 175th to 290th amino acid (Fig. 4C). The nature of the interactions driving such intermolecular contacts are primarily *π*-*π*, cation-*π*, and electrostatic, as these domains are enriched in aromatic and charged residues, including a significant number of glutamic (E) and aspartic (D) acids in TDP-43, lysines (K) and arginines (R) in HSP70, as well as tyrosines (Y) and phenylalanines (F) in segments of both sequences (please see Figs. 2B and 4A). Cation-*π* and *π*-*π* interactions are mostly formed between aromatic residues within this region of HSP70 (from the 250th to the 300th residue) and K, R, Y and F amino acids of TDP-43 within the two aforementioned domains of TDP-43 and its LCD. Nevertheless, the overall frequency of contacts between HSP70 and TDP-43 is lower than that of TPD-43 with GR_25_ and PR_25_ as shown in Fig. 2B—being the highest value of heterotypic contacts per residue 0.05% between TDP-43 and HSP70 whilst almost 4% between both arginine-rich peptides and TDP-43. Thus, the positioning of HSP70 at the interfaces is likely driven by its low effective valency with TDP-43 (and GR25; Fig. S2), with only a few specific regions interacting, such as the CTD and LCD of HSP70 and TDP-43, respectively among few others (Fig. 4C). This low effective valency with TDP-43 and GR25 may also contribute to reducing the condensate’s surface tension with the diluted phase, as previously shown for low-valency colloidal patchy particles forming multilayer condensates^166,167^.

**FIG. 4.**
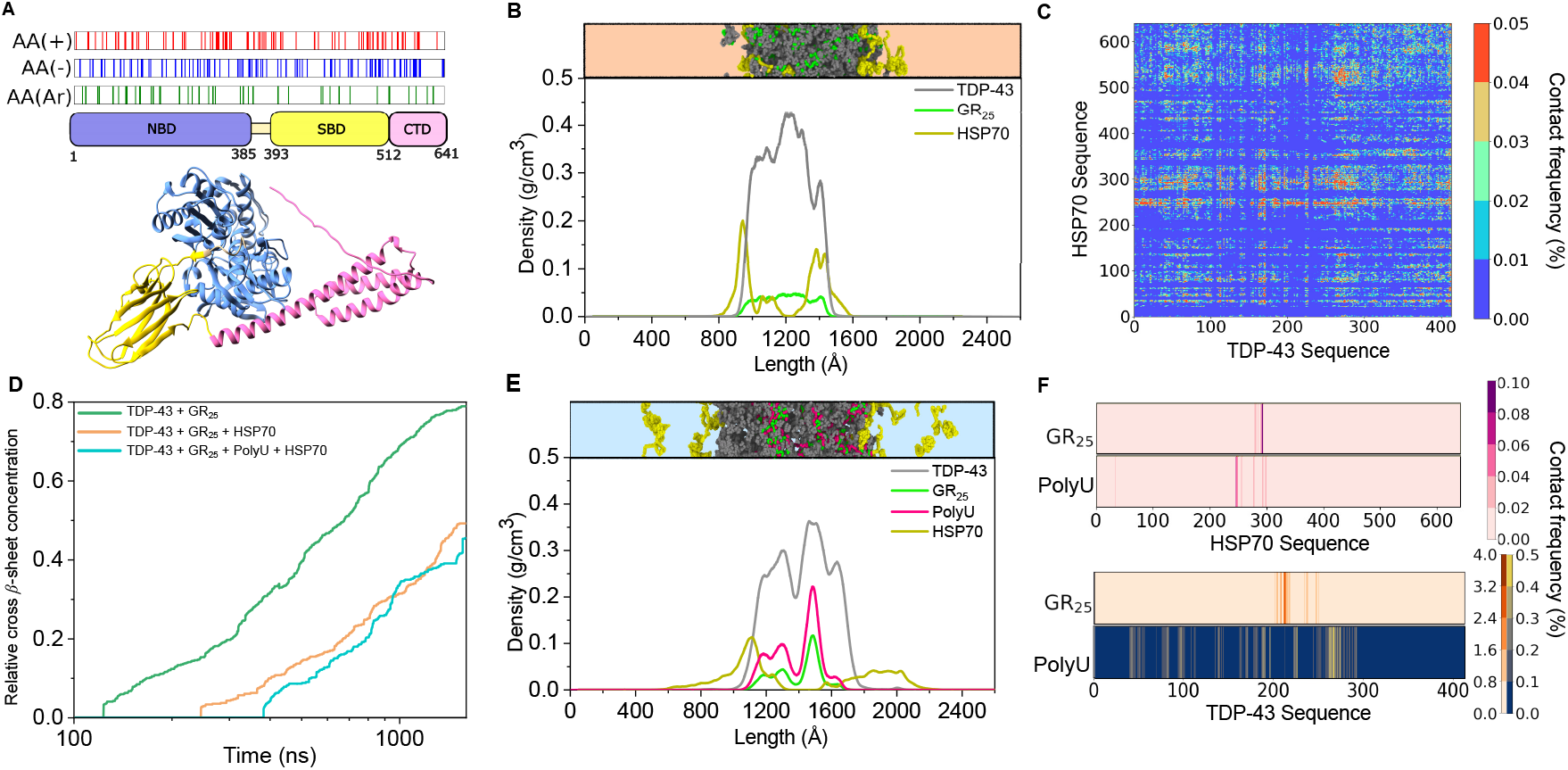
(A) Scheme of the HSP70 chaperone (1B Homo sapiens NP005337.2.) sequence displaying its different domains: a nucleotide binding domain (NBD) at the N-terminal, a substrate binding domain (SBD), and a C-terminal domain (CTD)^158^. The distribution of the positively and negatively charged amino acids as well as the aromatic residues across the sequence is shown above. (B) Density profile of a DC simulation at T=285K composed of TDP-43 (grey), HSP70 (khaki), and GR_25_ (green) at 1:0.16:1 molar ratio. The snapshot on top of the panel shows a representative configuration of the DC simulation. (C) Intermolecular contact frequency (in % of contacts per residue) between HSP70 and TDP-43 evaluated in the condensate shown in Panel B. (D) Time-evolution of the cross-*β*-sheet concentration (averaged for 5 independent trajectories and normalised by the maximum number of *β*-sheets which can be formed in the condensate) for different mixtures of TDP-43 condensates as indicated in the legend. (E) Density profile of a DC simulation at 285K composed of TDP-43 (grey), GR_25_ (green), polyU (pink) and HSP70 (khaki) with a molar stoichiometry of 1:1:0.16:0.16, respectively. Snapshot on top of the panel shows a representative configuration of the DC simulation. (F) Intermolecular contact frequency of polyU and GR_25_ with both HSP70 (Top panel) and TDP-43 (Bottom panel) in the condensate shown in Panel E.

We also perform non-equilibrium DC simulations to measure the kinetics of formation of inter-protein *β*-sheet assemblies in TDP-43/GR_25_/HSP70 condensates. We run five independent trajectories with different initial velocities and we measure the concentration of cross-*β*-sheet structures over time. In Fig. 4D, we compare the time-evolution of the average relative concentration of cross-*β*-sheets in condensates formed by TDP-43 and GR_25_ (green curve), and condensates of TDP-43/GR_25_/HSP70 (orange curve) with the same molar ratio as that shown in Fig. 4B. Remarkably, we find that the nucleation time for the formation of the first cross-*β*-sheet nucleus is significantly decelerated by a factor of 2 when the chaperone is present in the condensate—despite being at a relatively low concentration (TDP-43/HSP70 1:0.16 molar ratio). Such deceleration in ageing kinetics is similar to that found in presence of polyU strands of 250 nucleotides at the same molar ratio of 1:0.16 (Fig. 3C). However, whilst the impact of both HSP70 and polyU on ageing kinetics is alike, the molecular mechanism by which they regulate it seems to be radically different. PolyU outcompetes TDP-43 to interact with arginine-rich peptides, releasing contacts among the peptides and the TDP-43 RRM2 domain, and indirectly decreasing LCD-LCD interactions by favouring RRM2-LCD contacts. Moreover, direct interactions between polyU and the IDR1 of TDP-43 (adjacent to several LARKS) contribute to frustrating cross-*β*-sheet formation as well as RNA-RNA electrostatic self-repulsion which partially reduces condensate density and decreases the probability of TDP-43-LCD stacking. In contrast, HSP70 intervenes through a completely different mechanism in which TDP-43 LCD-LCD interactions seem to be precluded by competing heterotypic contacts with the CTD of HSP70. Moreover, the primarily localization of HSP70 towards the interface can contribute to hinder LCD-LCD stacking (substituted by CTD-LCD interactions) over the interfacial and near-interfacial regions, which as shown in Fig. 2D, enhance inter-protein *β*-sheet transitions compared to bulk condensate conditions.

Next, we investigate whether the addition of polyU modifies the internal organisation of TDP-43/GR_25_/HSP70 condensates. We perform DC simulations of a system with the same molar ratio as in Fig. 4B but including polyU strands of 250-nucleotides in the same molar ratio as HSP70 (e.g., TDP-43/GR_25_/HSP70/polyU: 1:1:0.16:0.16). In Fig. 4E, we show a snapshot of the condensate as well as the density profile across the long axis of the simulation box for each of the different species. Notably, whilst GR_25_ and polyU predominantly localise within the core of the condensate, HSP70 accumulates at the interface exhibiting low mixing with both RNA and GR_25_. TDP-43 recruits arginine-rich peptides to the core of the condensate due to their high-affinity binding through the RRM2 region. GR_25_ peptides additionally interact with polyU RNA through electrostatic-driven interactions. Meanwhile, HSP70 predominantly interacts with TDP-43 through CTD-LCD contacts, as well as through interactions between its NBD (mainly from the 250th to the 300th residue) and the NTD/RRM2 domains of TDP-43. The intermolecular contact maps between GR_25_ and polyU with both TDP-43 and HSP70 within the 4-component condensate are shown in Fig. 4F. We consistently find that GR_25_ predominantly contacts with the RRM2 domain of TDP-43 (rich in aromatic and negatively charged residues) and polyU strands interact with different segments across the TDP-43 sequence enriched in positively charged residues such as arginine and lysine, as shown in Fig. 2B. Strikingly, the contact frequency maps of GR_25_ and polyU with TDP-43 barely vary in presence of HSP70 (at this concentration) as shown in Figs. 2B, 3B, and 4F. Such contact network distribution is consistent with its known propensity to bind arginine-rich peptides and stabilise condensates in presence of nucleic acids^61^. In contrast to TDP-43, heterotypic contacts between HSP70 and both polyU and GR_25_ are scarce (Fig. 4F; Top panel). Specifically, GR_25_ interacts with a short segment of the HSP70 NBD region—rich in negatively charged residues (D_285_, E_283_, E_289_, D_292_)—and polyU RNA interacts with different short segments across the sequence containing lysines and arginines (K_246_, K_248_, K_250_, K_251_, K_257_, R_299_, R_301_). Such small extent of heterotypic contacts with both polyU and GR_25_ further contributes to primarily drive HSP70 localization towards the condensate interface. In that sense, HSP70 can act as a molecular buffer^168^, preventing excessive protein compaction and regulating condensate fluidity through specific interactions with hydrophobic protein domains as the TDP-43 LCD. Furthermore, the primary co-localisation of HSP70 at the interfaces may play a functional role in protecting the condensate from premature ageing initiated in these regions ^76,142^, as well as contributing to controlling its growth.

We now measure the formation of inter-protein *β*-sheets in TDP-43/GR_25_ condensates containing both HSP70 and polyU (Fig. 4D). The addition of RNA in TDP-43/GR_25_/HSP70 condensates further delays the emergence of cross-*β*-sheet nuclei. The average nucleation time (from 5 different independent trajectories) for the onset of cross-*β*-sheet growth is over 3 times faster in TDP-43/GR_25_ condensates (green curve) than in the 4-component system (blue curve). This result suggests that HSP70 can further decelerate ageing when RNA is present, consistent with its established role as an anti-aggregation chaperone^168^. On the one hand, polyU strands compete with TDP-43 for establishing LLPS-stabilising interactions with GR_25_, thus indirectly hindering LCD-LCD intermolecular contacts by promoting RRM2-LCD contacts in TDP-43, and polyU-IDR1 TDP-43 interactions which partially destabilize LCD-LCD high-density fluctuations due to RNA-RNA electrostatic self-repulsion. On the other hand, HSP70 at the interface directly competes with TDP-43 to form LCD-LCD intermolecular contacts through heterotypic CTD-LCD interactions. The synergy between the indirect mechanism by which RNA reduces the frequency of TDP-43 LCD-LCD contacts, and the direct interaction of HSP70 with the TDP-43 LCD, as well as the chaperone’s location at the interface, notably decelerates the emergence of cross-*β*-sheet structures in TDP-43 condensates.

### E. TDP-43 LCD intermolecular contact probability dictates the onset of cross-*β*-sheet nuclei formation

In previous sections, we have shown how the composition of TDP-43 condensates modulates the propensity of TDP-43 LARKS to undergo structural transitions into cross-*β*-sheet assemblies. Whilst the inclusion of GR_25_ peptides speed up ageing kinetics driven by inter-protein *β*-sheet transitions, both polyU and HSP-70 decelerate their emergence. To further understand the precise mechanism regulating such behaviour, we now analyse the frequency of TDP-43 intermolecular contacts in the two most different case scenarios presenting the earliest (Fig. 5A; TPD-43/GR_25_) and latest (Fig. 5B; TDP-43/GR_25_/polyU/HSP70) formation of cross-*β*-sheet nuclei. We observe that TDP-43 intermolecular contacts (in pre-aged liquid-like condensates) decrease almost 50% upon the addition of both HSP-70 and polyU. While the effective reduction of TDP-43 intermolecular contacts is partly due to steric hindrance from additional species within the condensate, we observe that different domains of TDP-43 exhibit varying changes in their contact frequencies. Intermolecular interactions among RRM2 domains of different protein replicas increase over 30% upon polyU and HSP-70 addition. However, the intermolecular contact probability between TDP-43 LCDs—those driving inter-protein *β*-sheet transitions— decrease by almost 30% (Fig. 5B). Such decrease can be mainly ascribed to CTD-LCD interactions between HSP70 and TDP-43 (which outcompete LCD-LCD TDP-43 contacts; Fig. 4C)), and polyU-IDR1 interactions (Fig. 3B). Overall, while the inclusion of additional species within the condensate contributes to the global reduction of TDP-43 intermolecular contacts, and thus ageing, we show here that is not necessarily always the case, since GR_25_ inclusion augments LCD-LCD TDP-43 contact probability, while polyU/HSP70 reduce LCD-LCD interactions and favour RRM2-RRM2 TDP-43 contacts—which do not drive inter-protein *β*-sheet transitions.

**FIG. 5.**
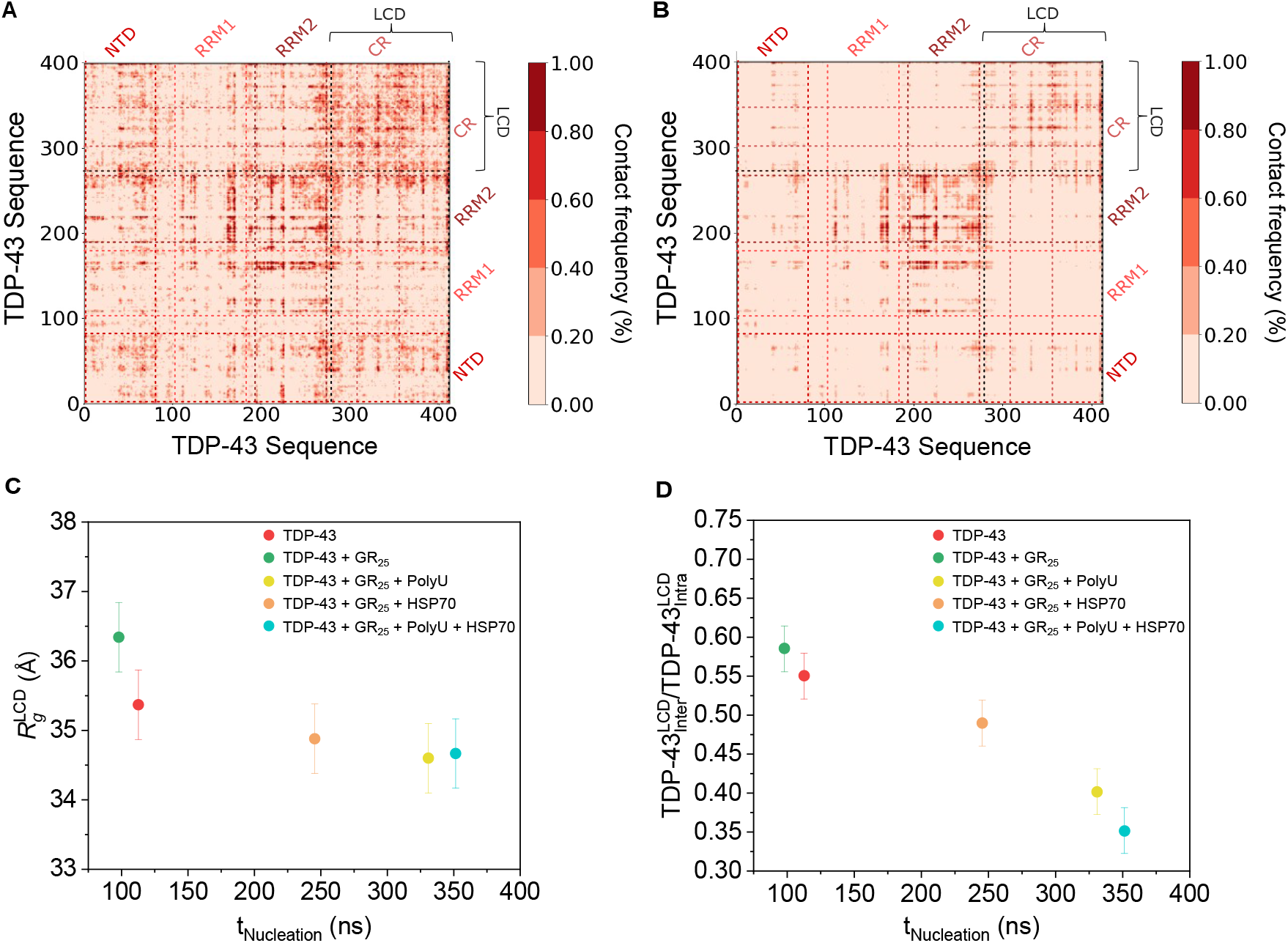
Map of TDP-43 intermolecular contacts (expressed in percentage per molecule residue) evaluated in TDP-43/GR_25_ condensates at 1:1 molar ratio (Panel A) and TDP-43/GR_25_/polyU/HSP70 condensates at 1:1:0.16:0.16 molar ratio (Panel B) at T = 285K at their corresponding condensate equilibrium density. Details on these calculations are provided in Section SV of the SM. (C) Radius of gyration (*R*_*g*_) of the low-complexity domain (LCD) of TDP-43 evaluated for different condensate mixtures as a function of their corresponding average nucleation time of cross-*β*-sheet clusters. (D) Ratio of intermolecular/intramolecular LCD contacts in TDP-43 for different condensate mixtures *vs*. their average nucleation time of cross-*β*-sheet formation.

Furthermore, since a decisive factor in TDP-43’s interprotein *β*-sheet transitions is how accessible are LCDs to interact with each other, we now evaluate the radius of gyration of the LCD of TDP-43 within condensates with the compositions previously studied. Remarkably, TDP-43/GR_25_ condensates display the largest *R*_*g*_ values in TDP-43 LCDs (Fig. 5C; green circle). Such more extended conformations are expected to facilitate inter-molecular contacts between different LCDs contributing to TDP-43 LCD favourable stacking and promoting inter-protein structural transitions. Indeed, TDP-43/GR_25_ condensates display the shortest timescale before the emergence of cross-*β*-sheet nuclei of all the studied systems. In contrast, multicomponent condensates— including both polyU RNA and HSP70 in addition to TDP-43 and GR_25_—promote TDP-43 LCD conformations which are moderately more compacted (Fig. 5C; cyan circle) and present the slowest nucleation time of cross-*β*-sheet formation (Fig. 4D). Hence, our simulations suggest a correlation between the LCD structural level of compaction—modulated by the presence of additional species—and the kinetics of cross-*β*-sheet formation (Fig. 5C). To further understand these differences in *R*_*g*_ among the different condensates, we also evaluate the ratio of inter-*versus* intramolecular LCD contacts (Fig. 5D). Higher ratios of intermolecular/intramolecular contacts are expected to favour LCD-LCD interactions triggering cross-*β*-sheet transitions. Fig. 5D reveals a strong correlation between higher inter/intra molecular contacts of the LCD and faster nucleation times of cross-*β*-sheet assemblies. This also indicates how larger *R*_*g*_ values of the LCD (i.e., more extended conformations) enable higher intermolecular contact frequencies, which in turn lead to faster ageing kinetics. Therefore, the way in which additional components to the scaffold protein—such as nucleic acids, peptides, or other proteins—regulate the accessibility of LARKS-containing LCDs determine the propensity to undergo cross-*β*-sheet transitions and progressive kinetic arrest over time.

## III. CONCLUSIONS

In this work, we investigate the impact of various biomolecules—such as arginine-rich peptides, RNA, and the HSP70 chaperone—on the stability, internal organisation, and ageing kinetics of condensates formed by TDP-43, a major RNA-binding protein that drives stress granule formation. We perform Molecular Dynamics simulations of the Mpipi-Recharged model^105^, a coarse-grained residue resolution force field which drastically improves the description of charge effects in biomolecular condensates, while maintaining the excellent predictions for low-complexity domains and multi-domains proteins of its predecessor Mpipi^169^, and still considering solvation effects implicitly for computational efficiency. We find that while the model moderately overestimates the experimental protein saturation concentration values of TDP-43 to undergo LLPS^54,110,111^(Fig. 1C), it closely reproduces *in vitro* experimental values of its radius of gyration under diluted conditions^109^ as well as atomistic modelling predictions^25^ (Fig. 1B). Furthermore, we benchmark the Mpipi-Recharged predictions of TDP-43 condensate stability as a function of different peptide di-repeat sequences, such as GP_25_, GR_25_, and PR_25_ (Fig. 2A) against *in vitro* experiments^50,52,108^. Remarkably, our model reproduces the re-entrant phase behaviour of TDP-43 condensates upon the addition of GR_25_ and PR_25_, capturing the precise peptide/protein molar ratio of condensate maximum stability^50,52,108^, and the monotonically decreasing stability after the inclusion of GP_25_^50^. Our simulations show that the main contacts enabling TDP-43 LLPS are interactions between the RRM2-RRM2, RRM1-RRM2, and LCD-LCD regions, mainly involving aromatic and charged residues that contribute to the condensate’s intermolecular connectivity. This is in agreement with previous experimental findings^124^ (Fig. 1E). However, in the presence of arginine-rich peptides, and at the stoichiometry that maximises the stability of the condensate (i.e., 1:1 molar ratio), the heterotypic contacts enhancing condensate stability are mostly electrostatic and cation-*π* interactions between GR_25_ (or PR_25_) and the RRM2 domain of TDP-43 which is rich in aromatic and negatively charged residues (Fig. 2B).

We perform non-equilibrium simulations using our ageing algorithm^82,139^ to investigate the formation of cross-*β*-sheet clusters in TDP-43 condensates over time. Interestingly, we discover that both pure TDP-43 and TDP-43/GR_25_ condensates exhibit faster nucleation of cross-*β*-sheet structures in condensates simulated in coexistence with the protein diluted phase than in condensate bulk conditions (Fig. 2D). These results suggest that inter-protein structural *β*-sheet transitions might be favoured at interfacial or near-interfacial regions in TDP-43 condensates as previously found for FUS^76,82^, *α*-synuclein^141^ and hnRNPA1^142^ condensates. Moreover, we find that the recruitment of arginine-rich peptides such as GR_25_ also accelerates the emergence of inter-protein *β*-sheet clusters as compared to pure TDP-43 systems (Fig. 2F). Hence, in addition to increasing their cohesion up to moderately high molar ratios, arginine-rich peptides also enhance their transition into aged kinetically trapped assemblies. The strong interaction that both GR_25_ and PR_25_ establish with the TDP-43 RRM2 region (Fig. 2B) partially releases RRM2-LCD contacts, thus indirectly increasing the frequency of LCD-LCD intermolecular interactions which trigger cross-*β*-sheet transitions.

We also determine the stability limits of TDP-43/GR_25_ condensates as a function of the concentration of polyU RNA. The addition of polyU moderately decreases the critical solution temperature of the condensates (Fig. 3A). PolyU predominantly co-localises at the condensate core strongly interacting with GR_25_ (Fig. 3B and Fig. S1) driven by attractive non-specific electrostatic interactions—in agreement with previous experimental and computational findings for other RNA-binding proteins including FUS^128,156^. The binding affinity of polyU is lower with TDP-43 than with GR_25_, only establishing significant interactions with its IDR1 domain and other different short segments along the sequence which are enriched in positively charged residues (Fig. 3B). Notably, the inclusion of polyU decelerates by twofold the nucleation time of cross-*β*-sheet assemblies compared to TDP-43/GR_25_ condensates (Fig. 3C), outcompeting with TDP-43 to interact with GR_25_ (Fig. S1) and overall reducing TDP-43 LCD intermolecular contacts. While polyU strands do not establish direct interactions with TDP-43 LARKS (Fig. 3B), they compete with TDP-43 to interact with GR_25_, releasing contacts between GR_25_ and the RRM2 domain of TDP-43, which consequently enable RRM2-LCD interactions hindering cross-*β*-sheet transitions. Moreover, direct interactions between polyU and the IDR1 of TDP-43, which has several adjacent LARKS, contribute to decelerating ageing kinetics according to our model. Additionally, the electrostatic self-repulsion among polyU strands at a moderately high concentration locally reduces the condensate density, thus hindering the frequency of TDP-43-LCD stacking promoting *β*-sheet transitions.

Furthermore, we introduce the chaperone HSP70 into TDP-43/GR_25_ condensates. HSP70 primarily concentrates at the condensate’s interface (Fig. 4B) establishing a low number of contacts with GR_25_ and several domains of TDP-43 sequence (i.e., NTD, RMM2 or LCD). The interactions between TDP-43 and HSP70 are mediated principally by *π*-*π*, cation-*π*, and electrostatic contacts, establishing the (HSP70) CTD and the (TDP-43) LCD the most relevant intermolecular interactions between both species (Fig. 4C). Such observation is consistent with previous work^165^ suggesting hydrophobic-like interactions to be critical for chaperones, in particular HSP70, in preventing protein aberrant aggregation. HSP70 decelerates the emergence of cross-*β*-sheet clusters by a factor of 2 (Fig. 4D), which is similar to the nucleation rate slow down induced through polyU inclusion. However, the mechanism by which HSP70 reduces inter-protein *β*-sheet transitions is through a direct competition of CTD-LCD contacts between HSP70 and TDP-43 *vs*. LCD-LCD interactions among TDP-43 proteins. The maximum deceleration in the formation of cross-*β*-sheet nuclei is achieved when both HSP70 and polyU are recruited within TDP-43/GR_25_ condensates (Fig. 4D). The synergy between the indirect mechanism by which RNA reduces the contact probability of TDP-43 LCD-LCD contacts, and the direct competition of HSP70 for TDP-43 LCD-LCD interactions delays the emergence of cross-*β*-sheet structures by a factor of 3 with respect to TDP-43/GR_25_ mixtures, which are those exhibiting fastest ageing kinetics. We note that the deceleration through polyU and HSP70 insertion can depend significantly on the stoichiometry of the mixture, as previously shown for FUS and hnRNPA1/polyU condensates^139^. Remarkably, even at low stoichimetric ratios of both polyU and HSP70 as studied here, their impact on ageing kinetics seems to be substantial.

Finally, we determine that the number of intermolecular contacts that TDP-43 homotypically establishes in the presence of GR_25_ (Fig. 5A) doubles the number upon addition of both polyU and HSP70 (Fig. 5B), despite the latter species being present at a much lower molar ratio (6 times less) than TDP-43 and GR_25_. While this reduction is in part due to steric hindrance from additional species within the condensate, we observe that intermolecular interactions between TDP-43 RRM2 domains increase in presence of polyU and HSP70 with respect to TDP-43/GR_25_ condensates—the system exhibiting fastest ageing kinetics. However, a 30% reduction in LCD-LCD contacts is achieved upon inclusion of polyU and HSP70. Furthermore, our simulations show a correlation between ageing kinetics and the ratio of intermolecular/intramolecular contacts of TDP-43 LCDs, which in turn also correlates with its degree of compaction evaluated through its radius of gyration (Figs. 5C and 5D). Condensate mixtures exhibiting faster nucleation times of cross-*β*-sheet transitions are scaffolded by TDP-43 proteins which in average possess more extended LCD conformations, and hence, higher probability of establishing LCD-LCD intermolecular contacts. The inclusion of different biomolecules to the scaffold protein—such as nucleic acids, peptides, or other proteins—regulates the accessibility of LCDs containing LARKS and determines the propensity to undergo cross-*β*-sheet transitions and progressive kinetic arrest. Altogether, our simulations provide molecular-level insights into the key interactions, mechanisms, and specific factors that might modulate the condensate phase behaviour of RNA-binding proteins, such as TDP-43, implicated in neurodegenerative diseases. Uncovering the microscopic pathways underlying aberrant liquid-to-solid transitions of condensates is crucial for developing potential therapeutic strategies to control condensate dynamics, given their implications in health and disease.

## Supporting information

SupportingInformation

## IV. ACKNOWLEDGEMENTS

A. F. acknowledges funding from the Ramon y Cajal fellowship (RYC2021-030937-I) and Spanish National Grant (PID2022-136919NA-C33). I. S.-B. acknowledges funding from Derek Brewer scholarship of Emmanuel College and EPSRC Doctoral Training Programme studentship, number EP/T517847/1, Ramon y Cajal fellowship (awarded to J.R.E.), as well as the UKRI EPSRC under the UK Government’s guarantee scheme (EP/Z002028/1), following successful evaluation by the ERC (Consolidator Grant awarded to R.C.G.) under the European Union’s Horizon Europe research and innovation programme. A.R acknowledges funding from PID2023-147156NB-I00 of the Spanish Ministry for Science, Innovation and Universities. R.C.-G. acknowledges funding from the European Research Council (ERC) under the European Union Horizon 2020 research and innovation programme (grant agreement 803326). A. R. T. acknowledges funding from the European Union Horizon 2020 research and innovation programme (grant agreement 803326 to R.C.-G.). J. R. E. acknowledges funding from Emmanuel College, the University of Cambridge, the Ramon y Cajal fellowship (RYC2021-030937-I), and the Spanish scientific plan and committee for research reference PID2022-136919NA-C33. M. M. C. acknowledges funding from the Ministerio de Ciencia e Innovación (Grant PID2022-136919NB-C32). This work has been performed using resources provided by the Cambridge Tier-2 system operated by the University of Cambridge Research Computing Service (http://www.hpc.cam.ac.uk) funded by EPSRC Tier-2 capital grant EP/P020259/1-CS170. This work has also been performed using resources provided by Archer2 (https://www.archer2.ac.uk/) funded by EPSRC Tier-2 capital grant EP/P020259/e829. The authors also thankfully acknowledge RES computational resources provided by Mare Nostrum 5 through the activity 2024-3-0001.

## V. DATA AVAILABILITY STATEMENT

We provide Supplementary Material and a Github repository that contains the LAMMPS input script as well as the configuration files of the different mixtures.

## Notes

https://github.com/Reshiiiii/TDP-43_Data_Scripts

### Competing Interest Statement

The authors have declared no competing interest.

