## SupportingInformation for "Compositional control of ageing kinetics in TDP-43 condensates"

### Simulation study of TDP-43 phase separation and fibrillization in presence of different biomolecules (Supplementary Material)

Nuria H. Espejo[+]

*Department of Physical-Chemistry Universidad Complutense de Madrid Av. Complutense s/n, Madrid 28040, Spain and Bayer CropScience, Alfred-Nobel-Straße 50, 40789 Monheim am Rhein, Germany*

Alejandro Feito[+], Antonio Rey, Alejandro Castro, and Jorge R. Espinosa[*]

*Department of Physical-Chemistry Universidad Complutense de Madrid Av. Complutense s/n, Madrid 28040, Spain*

Ignacio Sanchez-Burgos and Rosana Collepardo-Guevara

*Yusuf Hamied Department of Chemistry, University of Cambridge, Lensfield Road, Cambridge CB2 1EW, UK*

Adiran Garaizar

*Bayer CropScience, Alfred-Nobel-Straße 50, 40789 Monheim am Rhein, Germany*

Maria M. Conde

*Department of Chemical Engineering, Universidad Politécnica de Madrid, C/ Jose Gutierrez Abascal 2, Madrid 28006, Spain*

Andres R. Tejedor[†]

*Department of Physical-Chemistry Universidad Complutense de Madrid Av. Complutense s/n, Madrid 28040, Spain and Yusuf Hamied Department of Chemistry, University of Cambridge, Lensfield Road, Cambridge CB2 1EW, UK*

*+ These authors contributed equally*

(Dated: February 21, 2025)

*

†

#### SI. MODEL AND METHODS

In our simulations, we represent the proteins as coarse-grained polymers with one bead per amino acid. The potential energy of the Mpipi-Recharged coarse-grained force field is defined as follows:

$$E = E_{\text{Bonds}} + E_{\text{Electrostatic}} + E_{\text{Hydrophobic}}. \tag{S1}$$

Bonded interactions are modelled by a harmonic potential

$$E_{\text{Bonds}} = \sum k(r_i - r_0)^2, \tag{S2}$$

where $k = 9.6\text{kJ/mol} \cdot \text{Å}^{-2}$ and the equilibrium bond length is $r_0 = 3.81$ Å between bonded amino acid beads.

Electrostatic interactions ($E_{\text{Electrostatic}}$) among charged amino acids are given by the Yukawa potential expressed as

$$E_{\text{Electrostatic}} = \sum_i \sum_{j<i} \frac{A_{ij}}{r} e^{-r\kappa}, \tag{S3}$$

where $A_{ij}$ controls the interaction between a pair of amino acids and $\kappa$ modulates the salt concentration in an explicit way. The Debye length $\kappa$ depends on the salt concentration as

$$\kappa = \sqrt{8\pi B c_s}, \tag{S4}$$

where $B(\epsilon_r) = e^2/4\pi k_B T \epsilon_0 \epsilon_r$ is the Bjerrum length, and the relative dielectric permittivity depends on the temperature according to this empirical law

$$\epsilon_r(T) = \frac{5321}{T} + 233.76 - 0.9297T + 1.41710^{-3}T^2 - 8.29210^{-7}T^3. \tag{S5}$$

Hydrophobic interactions are parameterised by the Wang-Frenkel potential given by

$$E_{\text{Hydrophobic}} = \sum_i \sum_{j<i} \epsilon_{ij} \alpha \left( \left[\frac{\sigma_{ij}}{r}\right]^{2\mu} - 1 \right) \left( \left[\frac{r_c}{r}\right]^{2\mu} - 1 \right)^{2\nu_{ij}} \tag{S6}$$

with

$$\alpha = 2\nu \left(\frac{r_c}{\sigma}\right)^{2\mu} \left[ \frac{1 + 2\nu_{ij}}{2\nu_{ij}\left(\left(\frac{r_c}{\sigma}\right)^{2\mu} - 1\right)} \right]^{2\nu_{ij}+1}, \tag{S7}$$

where the excluded volume of the different residues is given by $\sigma_{ij}$, $r$ is the distance between the $ij$ particles, $\epsilon_{ij}$ is the energy interaction parameter for every pair of residues, $r_c = 3\sigma_{ij}$

is the cut-off of the potential between those amino acids and $\mu$=1 and $\nu_{ij}$ are terms that are involved in the shape of the potential. The values of $\sigma_{ij}$ and $\epsilon_{ij}$ are precisely parametrised for each interaction [1, 2]. To account for buried interactions, the energy interaction between two amino acids in a globular is $0.7\epsilon_{ij}$, and $\sqrt{0.70}\epsilon_{ij}$ between one in a structured region and another in a disordered segment. We employ a cut-off of $3\sigma_{ij}$ for the hydrophobic interactions, and 3.5 nm for the electrostatic ones [3].

#### SII. PREPARATION OF THE SYSTEMS AND SIMULATION DETAILS

All simulations were conducted using the LAMMPS software [4, 5]. All simulations are performed at physiological NaCl concentration, 150 mM. We also turn off the interactions between particles that are part of the same globular structured domain (i.e., within the same rigid body) using the LAMMPS package RIGID.

The simulations in the NVT ensemble use a Langevin thermostat [6] with a relaxation time of 5 ps and a time step of 10 fs for the Verlet integration.

For calculating the radius of gyration we run NVT simulation of one single TDP-43 replica in a cubic box of 400 Angstrom length for $\sim$ 10 ns upon equilibration of $\sim$0.1 ns. The configurations for the DC simulations are generated by placing 100 protein replicas in a slab box with $\sim$20x20 nm$^2$ of section and 140 nm in the long side for a resulting density of $\sim$0.13 g/cm$^3$. The production run was of the order of $\sim$1$\mu$s after reaching equilibrium.

We run NpT simulations to calculate the critical temperature of multicomponent systems, using a Langevin thermostat with a relaxation time of 5 ps and a time step of 10 fs for the Verlet integration and a Berendsen barostat [7] with a relaxation of 5 ps. We determined the critical temperature by considering a system density below $\sim$ 0.1 g/cm$^3$ as critical. The configurations are generated by placing 100 protein replicas of TDP-43 and the corresponding replicas of the other components depending on that system composition in bulk conditions (cubic box).

Simulations using the dynamic algorithm (see SVII), are performed in the NVT ensemble employing a Langevin thermostat with a relaxation time of 5 ps and a time step of 10 fs for the Verlet integration, using the same method of DC or bulk conditions previously explained, but with 64 protein replicas of TDP-43 and the corresponding replicas of the other components, depending on the composition of the specific system.

#### SIII. SEQUENCE OF TDP-43 AND PDB OF THE STRUCTURED DOMAINS AND SEQUENCE OF HSP70

TDP-43:

```
MSEYIRVTEDENDEPIEIPSEDDGTVLLSTVTAQFPGACGLRYRNPVSQCMRGVRLVEGILHAPDAGWGNLVYVV
NYPKDNKRKMDETDASSAVKVKRAVQKTSDLIVLGLPWKTTEQDLKEYFSTFGEVLMVQVKKDLKTGHSKGFGFV
RFTEYETQVKVMSQRHMIDGRWCDCKLPNSKQSQDEPLRSRKVFVGRCTEDMTEDELREFFSQYGDVMDVFIPKP
FRAFAFVTFADDQIAQSLCGEDLIIKGISVHISNAEPKHNSNRQLERSGRFGGNPGGFGNQGGFGNSRGGGAGLG
NNQGSNMGGGMNFGAFSINPAMMAAAQAALQSSWGMMGMLASQQNQSGPSGNNQNQGNMQREPNQAFGSGNNSYS
GSNSGAAIGWGSASNAGSGSGFNGGFGSSMDSKSSGWGM
```

The following Protein Data Bank (PDB) codes were used to build the globular structured domains of TDP-43 residues: 5MDI for NTD (2-38, 40-49, 51-79) [8]; 2CQG for RRM1 (103-179) [9], 1WF0 for RRM2 (193-267) [10] and 2N2C for CR (307-349) [11]). For the CR we increase the $\epsilon_{ij}$ between this region itself a 10%. The rest of the intervals in the sequence are maintained as flexible parts.

HSP70:

```
MAKAAAIGIDLGTTYSCVGVFQHGKVEIIANDQGNRTTPSYVAFTDTERLIGDAAKNQVALNPQNTVFDAKRLIG
RKFGDPVVQSDMKHWPFQVINDGDKPKVQVSYKGETKAFYPEEISSMVLTKMKEIAEAYLGYPVTNAVITVPAYF
NDSQRQATKDAGVIAGLNVLRIINEPTAAAIAYGLDRTGKGERNVLIFDLGGGTFDVSILTIDDGIFEVKATAGD
THLGGEDFDNRLVNHFVEEFKRKHKKDISQNKRAVRRLRTACERAKRTLSSSTQASLEIDSLFEGIDFYTSITRA
RFEELCSDLFRSTLEPVEKALRDAKLDKAQIHDLVLVGGSTRIPKVQKLLQDFFNGRDLNKSINPDEAVAYGAAV
QAAILMGDKSENVQDLLLLDVAPLSLGLETAGGVMTALIKRNSTIPTKQTQIFTTYSDNQPGVLIQVYEGERAMT
KDNNLLGRFELSGIPPAPRGVPQIEVTFDIDANGILNVTATDKSTGKANKITITNDKGRLSKEEIERMVQEAEKY
KAEDEVQRERVSAKNALESYAFNMKSAVEDEGLKGKISEADKKKVLDKCQEVISWLDANTLAEKDEFEHKRKELE
QVCNPIISGLYQGAGGPGPGGFGAQGPKGGSGSGPTIEEVD
```

The structure used in this study is the AlphaFold prediction AF-P0DMV8-F1-v4, and the globular structured domains were built regarding the regions scoring higher than 90%, in the per-residue model confidence score (pLDDT). The residues forming these regions are:

- NTD and SBD domains: 11-14, 19-26, 29-32, 46-48, 53-61, 66-71, 73-78, 84-92, 97-101, 104-111, 114-118, 119-140, 145-150, 155-169, 172-178, 179-187, 196-204, 209-217, 220-229, 233-254, 260-278, encompassed in the same globular structured domain, and 283-292, 295-302, 303-333, 337-358, 371-382, 389-396, 401-405, 409-414, 422-430, 438-

444, 449-461, 474-480, 486-492, 498-503 in a different one.

- CTD domain: 505-509, 511-559 and 563-584. Separated in 3 different globular domains.

The rest of the intervals in the sequence are maintained as flexible parts.

#### SIV. CALCULATING RADIUS OF GYRATION

We analyse the single-protein conformational ensemble of TDP-43's sequence using the radius of gyration calculated as

$$R_g^2 = \frac{1}{M}\sum_i m_i(r_i - r_{cm})^2. \tag{S8}$$

where $M$ is the number of amino acids in the chain, $m_i$ is the mass of the amino acid, $r_i$ is the position of the amino acid and $r_{cm}$ is the position of the center of mass of the protein. For this purpose, we compute the distribution of the radius of gyration ($R_g$) of the protein chains in the phase. To measure $R_g$ in the diluted phase, we performed NVT simulations with a single protein at a temperature approximately $0.95T_c$ of the critical temperature. Upon reaching equilibrium, we compute the $R_g$ histograms along the simulation.

#### SV. DIRECT COEXISTENCE TECHNIQUE

Using direct co-existence (DC) simulations [12–15], we determine the phase diagram for TDP-43. This method involves simulating the two coexisting phases within the same simulation box. In our approach, we arrange a high-density protein liquid alongside a very low-density counterpart. To accommodate the diverse densities, we employ an elongated simulation box, therefore allowing for the formation of both coexisting phases forming an interface perpendicular to the long direction of the slab. After equilibrium is achieved, we measure the equilibrium coexisting densities of both phases along the elongated side of the box. This process is repeated atrious temperatures until thethe critical point is reached.To mitigate finite system-size effects near the critical point, we calculate the critical temperature ($T_c$) and density ($\rho_c$) using the law of critical exponents and rectilinear diameters [16] given by

$$(\rho_l - \rho_v)^{\alpha} = s_1\left(1 - \frac{T}{T_c}\right) \tag{S9}$$

and

$$\frac{\rho_l + \rho_v}{2} = \rho_c + s_2(T_c - T), \tag{S10}$$

where the notation $\rho_l$ and $\rho_v$ are the densities of the condensed and diluted phases, respectively. Moreover, $s_1$ and $s_2$ are fitting parameters, while the critical exponent, $\alpha = 3.06$ for the three-dimensional Ising model [16].

#### SVI. CALCULATING CONTACT MAPS

Computation of intermolecular and intramolecular contact maps within protein condensates is calculated from DC trajectories. Contacts are determined across all systems at temperatures of approximately $0.95T_c$, where $T_c$ corresponds to the critical temperature of each system. Typically, molecular contacts are identified based on a distance criterion, with the assumption that the relative frequency of contact map occurrences (rather than absolute frequency) remains generally unaffected by the selected cut-off distance used in calculations, assuming the cut-off values are reasonable. In particular, we use a criterion that depends on the specific excluded of each amino acid, adopting a sequence-dependent cut-off distance equivalent to $1.2\sigma_{ij}$, where $\sigma_{ij}$ represents the mean excluded volume of the respective ith and jth amino acids [17]. Given that the minimum of the potential used lies at approximately $2^{1/6}\sigma_{ij} \approx 1.122\sigma_{ij}$, we set the cut-off distance slightly beyond this point, at $1.2\sigma_{ij}$, to ensure significant binding.

For those molecules with periodic sequence (i.e. PolyU and arginine dipeptide repeats), we have computed the average number of contacts of all the residues along the sequence and provided the result in a map. In Fig. S1 we plot the contact maps of PolyU with $GR_{25}$ (top) and $PR_{25}$ (bottom). We also provide the intermolecular contact maps of $GR_{25}$ (Fig. S2) and the contact map of HSP-70 with TDP-43 in the system containing $GR_{25}$ (Fig. S3).

#### SVII. DYNAMIC ALGORITHM FOR AGEING

To dynamically mimic the structural disorder-to-order transitions of TDP-43 LARKS during ageing, we employ a time and local dependent coarse-grained algorithm developed by us [18, 19]. Our dynamic algorithm enable effective structural transitions from disorder to form inter-protein $\beta$-sheets by increasing the binding energies and stiffness of the target

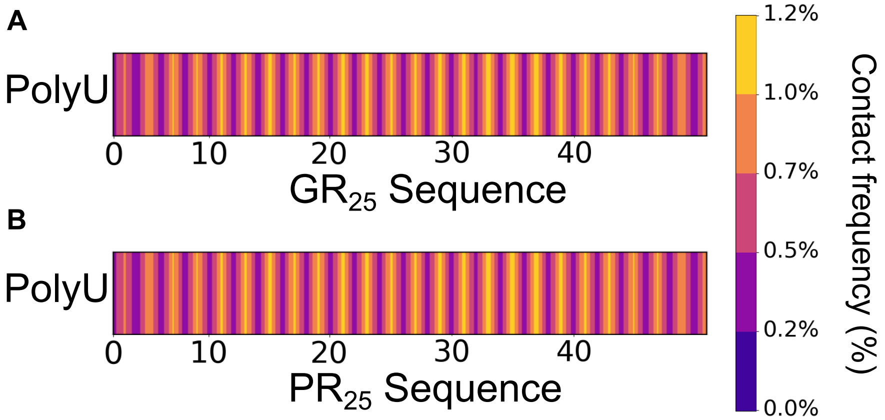


**FIG. S1:** (A) Intermolecular frequency contact maps for $GR_{25}$ dipeptide and PolyU for the system with TDP-43, $GR_{25}$ and PolyU. (B) Intermolecular frequency contact maps for $PR_{25}$ dipeptide and PolyU for the system with TDP-43, $PR_{25}$ and PolyU. The contacts are represented as contact frequencies in percentage.

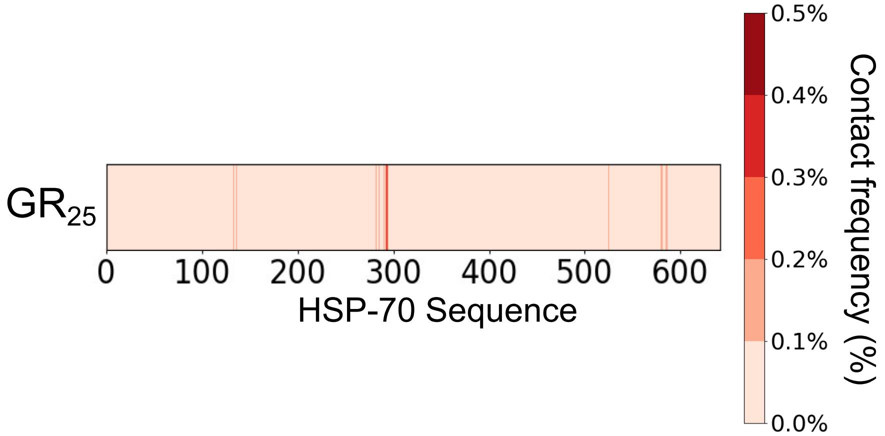


**FIG. S2:** Intermolecular frequency contact maps for HSP70 protein and $GR_25$ for the system with TDP-43, $GR_{25}$ and HSP70. The contacts are represented as contact frequency in percentage. The frequencies have been represented in red tones.

LARKS. In particular, in this work we considered the following LARKS: $_{312}$NFGAFS$_{317}$ as LARKS 1; $_{333}$SNGMMGMLASQ$_{343}$ as LARKS 2; and $_{396}$GFNGGFG$_{402}$ as LARKS 3. Every Every 100 simulation time steps, the algorithm evaluates the distance between LARKS and triggers the disorder-to-order transition following these conditions: (1) two LARKS of the same type are within a cut-off distance of 13.5 Å, and (2) both LARKS are surrounded by at least other 2 LARKS within a cut-off distance of 13.5 Å. Under this assumptions,

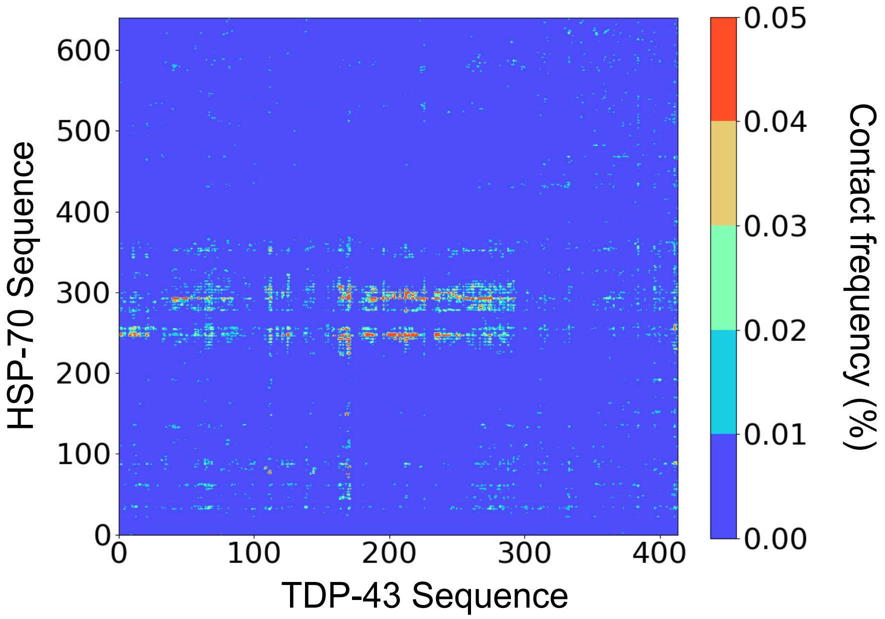


**FIG. S3:** Intermolecular frequency contact maps for TDP-43 protein and HSP70 chaperone for the system with TDP-43, $GR_{25}$, poly-U and HSP70. The contacts are represented as contact frequency in percentage.

the algorithm changes the identity of the LARKS residues to increase the LARKS-LARKS interaction and they are assigned a mass and $\sigma$ to the total mass constant. The specific parameters corresponding to each LARKS after the transition occurs as well as the residue used to estimate the distance between LARKS are provided in Table S1. To carry out these simulations, we use the REACTION [20] package of LAMMPS, which allows us to change the identity and thus, the interaction of the system components on the fly.

Furthermore, to capture the local increase in rigidity linked to a $\beta$-sheet structural transition, we incorporate an angular term into the total energy (Eq. S11), expressed as

$$E_{\text{Angles}} = \sum_{\text{Angles}} k_{\text{ang}}(\theta - \theta_0)^2, \tag{S11}$$

where $\theta_0 = 180°$ and $k_{\text{ang}} = 5\,\text{kcal}\cdot\text{mol}^{-1}\cdot\text{rad}^{-2}$ for the structured LARKS in an inter-protein $\beta$-sheet.

Finally, it is important to note that when LARKS 1 forms a $\beta$-sheet structure, we also include the segments $_{300}$GNNQGSN$_{306}$ and $_{328}$AALQSS$_{333}$, but increasing the interaction energy only with themselves to simulate a parallel $\beta$-sheet as reported experimentally [21]. In that respect, since $S_{333}$ is present in both LARKS 1 and LARKS 2, only one of these

| | LARKS 1 | LARKS 2 | LARKS 3 |
|---|---|---|---|
| Reactive amino acid | $G_{315}$ | $S_{333}$ | $G_{399}$ |
| $m_{\rm ordered}$ (g mol$^{-1}$) | 95.94 | 103.95 | 107.31 |
| $\epsilon_{\rm ordered}$ (kcal mol$^{-1}$) | 1.23 | 1.32 | 1.17 |
| $\sigma_{\rm ordered}$ (Å) | 5.66 | 5.59 | 5.96 |

**TABLE S1:** Set of parameters employed for residues belonging to structured inter-peptide $\beta$-sheet motifs in TDP-43. LARKS 1, LARKS 2 and LARKS 3 correspond to the $_{312}$NFGAFS$_{317}$, $_{333}$SNGMMGMLASQ$_{343}$ and $_{396}$GFNGGFG$_{402}$ sequences respectively. The reactive amino acid indicates the residue used by the ageing algorithm to evaluate LARKS high-density fluctuations. $m_{\rm ordered}$, $\epsilon_{\rm ordered}$, and $\sigma_{\rm ordered}$ refer to the mass, interaction energy, and molecular diameter respectively of the residues composing the structured LARKS of each sequence. Please note that in our model, after the structural transition takes place, the different residues of a given sequence are replaced by an average mutated residue that conserves the volume and mass of the unstructured LARKS.

processes can occur within the same replica of TDP-43.

#### SVIII. MEAN SQUARED DISPLACEMENT CALCULATION

We calculate the mean squared displacement (MSD) of the centre of mass defined as

$$MSD(t) = \left\langle (\mathbf{r}_{CM}(t) - \mathbf{r}_{CM}(0))^2 \right\rangle , \tag{S12}$$

where $\mathbf{r}_{CM}(t)$ is the position centre of mass at time $t$, and $\mathbf{r}_{CM}(0)$ is the initial position of the centre of mass. For this calculation, we used a multiple-tau correlator [22] to improve the average that goes over all protein replicas in the system.

[10] F. He, Y. Muto, M. Inoue, T. Kigawa, M. Shirouzu, T. Terada, and S. Yokoyama, Solution structure of rrm domain in tar dna-binding protein-43, PDB ID: 1WF0 (2004).

[16] J. S. Rowlinson and B. Widom, *Molecular theory of capillarity* (Courier Corporation, 2013).
